# Tic-related behaviors in *Celsr3* mutant mice are contributed by alterations of striatal D_3_ dopamine receptors

**DOI:** 10.1101/2023.12.07.570669

**Authors:** Roberto Cadeddu, Caterina Branca, Giulia Braccagni, Teresa Musci, Ignazio S Piras, Collin J Anderson, Mario R Capecchi, Matthew J Huentelman, Philip J Moos, Marco Bortolato

## Abstract

The gene *CELSR3* (Cadherin EGF LAG Seven-pass-G-type Receptor 3) has been recently recognized as a high-confidence risk factor for Tourette syndrome (TS). Additionally, *Celsr3* mutant mice have been reported to exhibit TS-related behaviors and increased dopamine release in the striatum. Building on these findings, we further characterized the neurobehavioral and molecular profile of *Celsr3* mutant mice to understand better the biological mechanisms connecting the deficiency of this gene and TS-related phenotypes. Our analyses confirmed that *Celsr3* mutant mice displayed grooming stereotypies and tic-like jerks, as well as sensorimotor gating deficits, which were opposed by TS therapies. Spatial transcriptomic analyses revealed widespread extracellular matrix abnormalities in the striatum of *Celsr3* mutants. Single-nucleus transcriptomics also showed significant upregulation of the *Drd3* gene, encoding the dopamine D_3_ receptor, in striosomal D_1_-positive neurons. *In situ* hybridization and immunofluorescence confirmed dysregulated D_3_ receptor expression, with lower levels in presynaptic striatal fibers and higher levels in striatal D_1_-positive neurons. Activating and blocking D_3_ receptors amplified or decreased tic-like jerks and stereotypies *in Celsr3*-deficient mice, respectively. These findings suggest that modifications of D_3_ receptor distribution contribute to the tic-like responses associated with *Celsr3* deficiency.

## INTRODUCTION

Tics are partially involuntary, highly repetitive movements or vocalizations with variable intensity and complexity. They are often a significant source of disability, negatively affecting social and emotional adjustment, educational attainment, and overall quality of life^1–5^. The most debilitating tic disorder is Tourette syndrome (TS), a neurodevelopmental disorder characterized by multiple motor tics and at least one vocal tic for over a year^6^. TS is estimated to affect approximately 0.5% of children^7^ and is notably more prevalent in boys, with a male-to-female ratio of ∼3:1^8^. The typical course of TS features a clinical onset around six years, followed by a progressive exacerbation of tic severity and a gradual amelioration after puberty^9, 10^. The trajectory of TS is also influenced by comorbid psychiatric conditions such as obsessive-compulsive disorder (OCD), attention-deficit hyperactivity disorder (ADHD), anxiety, and depression^11–14^. The primary drug treatment for TS involves the use of the D_2_ receptor antagonist haloperidol or the α_2_ adrenergic receptor agonist clonidine ^15^; however, the efficacy of these treatments is inconsistent, and their utility is hampered by significant adverse effects^16, 17^, underscoring the critical need to identify novel therapeutic targets.

The pathophysiology of tics is associated with disruptions in the cortico-striatal-thalamo-cortical (CSTC) circuitry^18^. Specifically, tics are posited to arise from abnormal disinhibition in focal areas of the dorsal striatum^19^, contributed by dopaminergic overstimulation^20-22^ and the loss of specific classes of interneurons^23-25^. Despite this evidence, the molecular underpinnings of TS remain largely unclear. Given the strong genetic predisposition of TS and other tic disorders, genetic studies are instrumental in uncovering their molecular foundations. Recent research on *de novo* mutations has identified *CELSR3* (cadherin epidermal growth factor laminin G seven-pass G-type receptor 3) as one of the first high-confidence risk genes for TS^26-28^. CELSR3 encodes a membrane receptor in the cadherin superfamily, which contains seven transmembrane domains and a large ectodomain interacting with neighboring cells^29^. This molecule is broadly expressed in the CNS, where it regulates the formation of neural pathways and basal ganglia^30-35^.

To examine the phenotypic effects of *CELSR3* mutations, a recent study developed mice harboring human-like *Celsr3* mutations and found that these mutants exhibited behavioral changes related to TS, including increased grooming stereotypies and deficits in prepulse inhibition (PPI)^36^, a process commonly impaired in TS patients^37, 38^. Despite these abnormal phenotypes, the mutants showed no obvious anatomical defects or interneuron loss but instead displayed increased dopamine release in the striatum^36^.

To investigate the molecular underpinnings of the tic-associated phenotypes linked to CELSR3 mutations, here we analyzed the transcriptomic profile of the striatum of a different line of *Celsr3*-deficient mice generated by the insertion of a neomycin resistance cassette^39, 40^. Since homozygous knockout mice for this gene die shortly after birth^34^, our analysis focused on heterozygous (HZ) and wild-type (WT) control mice. The HZ genotype is particularly relevant because *CELSR3 de novo* mutations, which typically affect only one allele, have been identified in individuals with TS^20–22^. After confirming the face and predictive validity of this mouse model for TS and its comorbid disorders (see Suppl. Fig.1), we conducted bulk, spatial, and single-nucleus transcriptomic analyses of the striata from *Celsr3*-deficient mice to gain insight into the mechanisms underlying the emergence of TS-related phenotypes resulting from the deficiency of this protocadherin. Our results point to a previously unknown role of dopamine D_3_ receptors, which appear to play an unexpected role in the development of certain tic-related abnormalities in these mutants.

## RESULTS

The following descriptions provide a summary of the results. For a comprehensive account and all statistical details, please refer to Supplemental Results.

### *Celsr3* HZ Mice Display TS-Related Behavioral Alterations

We first examined the temporal trajectory of the behavioral responses of *Celsr3* HZ mice compared to WT littermates. No differences were noted in motor responses or ultrasonic vocalizations among male and female newborns of both genotypes during the first week of life (Suppl. Fig.2A-C). To investigate TS-related behaviors, we assessed tic-like jerks and stereotypies, as well as prepulse inhibition (PPI) of the acoustic startle, which is typically defective in TS patients ^37^. This evaluation was conducted in male and female HZ and WT littermates, using two separate cohorts aged 21 (immediately after weaning) and 42 days (juvenile stage) to avoid potential stress carryover effects. At 21 days, both male and female *Celsr3* HZ mice exhibited increased grooming relative to WT mice (Fig.1A), while jerks, chewing stereotypies, and eye blinking showed no significant differences between genotypes (Figs.1B-D). However, female HZ mice displayed decreased digging compared to WT littermates (Fig.1E). PPI deficits were evident across all loudness levels in male, but not female, HZ mice, despite no differences in acoustic startle amplitude (Figs.1F-G). At 42 days, grooming increased further in male and female HZ mice, with more pronounced effects in males (Fig.1H). Additionally, male *Celsr3* mutants exhibited more head and body jerks than WT and female HZ mice (Fig.1I), while chewing, blinking, and digging behaviors remained unchanged across groups (Figs.1J-L). Female mice showed a lower startle response than males, with male HZ mice displaying reduced PPI compared to WT males (Figs.1M-N). Overall, these tests indicated a high level of face validity for *Celsr3* mutant mice with respect to key deficits observed in TS patients. To verify whether these responses were accompanied by other behaviors related to comorbid diagnoses, we tested a separate cohort of juvenile (42-through 50-day-old) mice on a series of tests aimed at capturing complementary dimensions of behavioral reactivity (Suppl. Fig.3). *Celsr3* HZ mice displayed increased locomotion and preference for the central area of an open field, irrespective of sex (Suppl. Figs.3A-B). Unlike females, male HZ mice displayed deficits in familiar object recognition (Suppl. Fig.3C). Finally, no differences were detected in the marble-burying task, the elevated plus maze, or the suspended wire-beam bridge, which capture compulsive, anxiety-like, and impulsive responses (Suppl. Figs.3D-H).

**Fig. 1.**
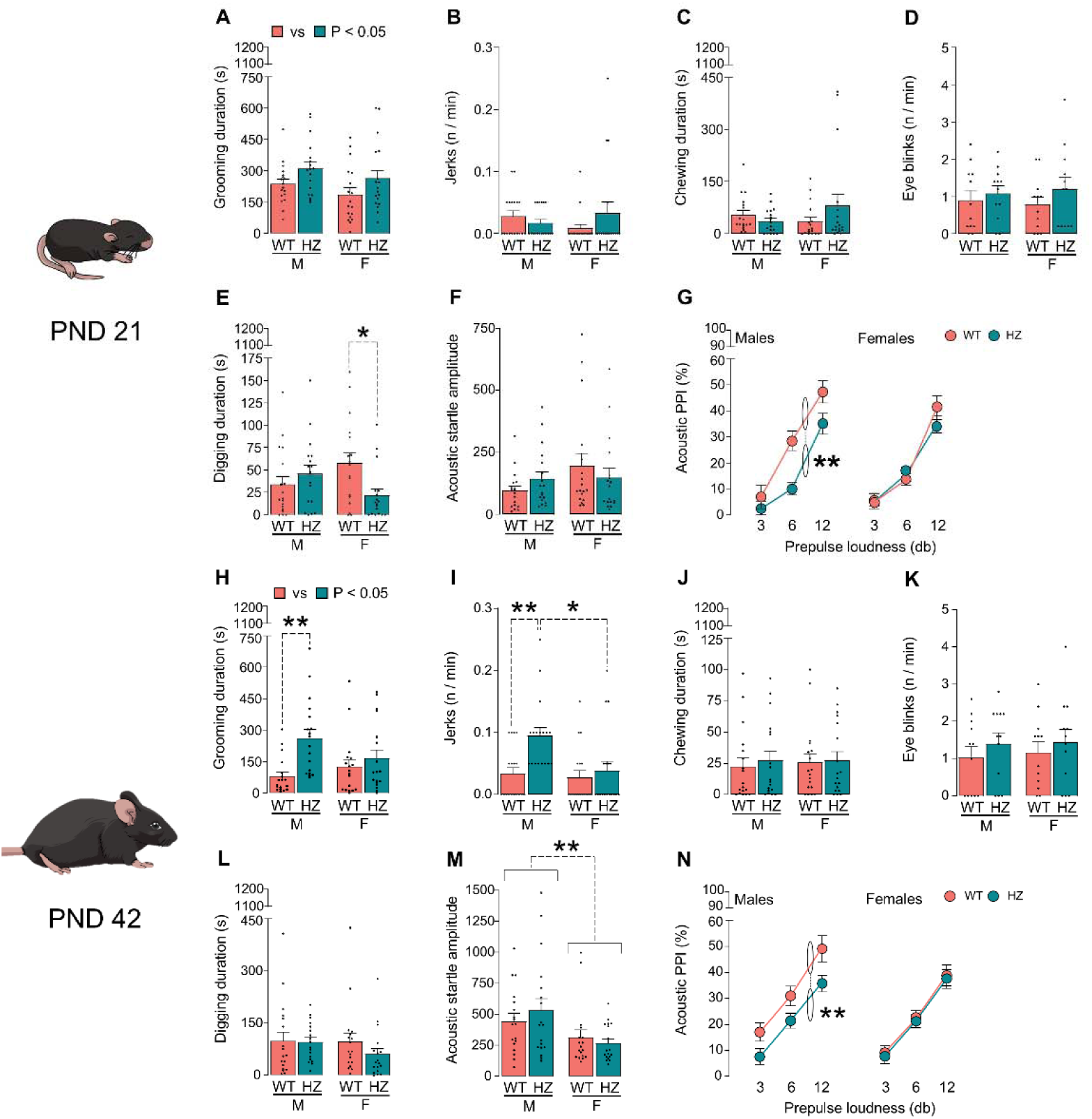
Adolescent and juvenile *Celsr3* heterozygous (HZ) mice show spontaneous behavioral alterations related to Tourette syndrome (TS). (A-G) Male and female *Celsr3* HZ mice were tested on postnatal day 21 (PND 21) for spontaneous TS-related behavioral alterations in comparison with wild-type (WT) littermates. *Celsr3* mutant mice displayed a significant increase in spontaneous grooming (A), but not differences in the frequency of body and head jerks (B), chewing stereotypies (C), and eyeblink (D). Female HZ mice displayed a significant reduction in the overall duration of digging behavior (E). While no difference in the amplitude of the acoustic startle reflex was found (F), *Celsr3* HZ males, but not females, displayed significant prepulse inhibition (PPI) deficits (G). (H-N) Spontaneous behaviors were also assessed in a separate group of juvenile mice (PND 42). Grooming duration (H) was increased in HZ mice, and the effects were particularly marked for males. *Celsr3* HZ males also showed a significant increase in the frequency of jerks compared to male WT and female HZ mice (I). Chewing (J), eye-blinking (K), and digging behaviors (L) were similar for all groups. Irrespective of genotype, males exhibited a significant increase in acoustic startle amplitude compared with females (M). However, only *Celsr3* HZ males displayed significant PPI deficits irrespective of the prepulse loudness (N). Data in panels G and N are presented as mean ± SEM for each sex and were analyzed by three-way ANOVA (for sex, genotype, and prepulse loudness). All other data are reported as mean ± SEM with individual data points and were analyzed by two-way ANOVA for sex and genotype (n=18/group, except for n=12/group for panels D and K). WT mice are indicated in coral, and HZ mice in teal. *, P<0.05; **, P<0.01 for comparisons indicated by brackets. For further details, see text and Suppl. Results.

**Fig. 2.**
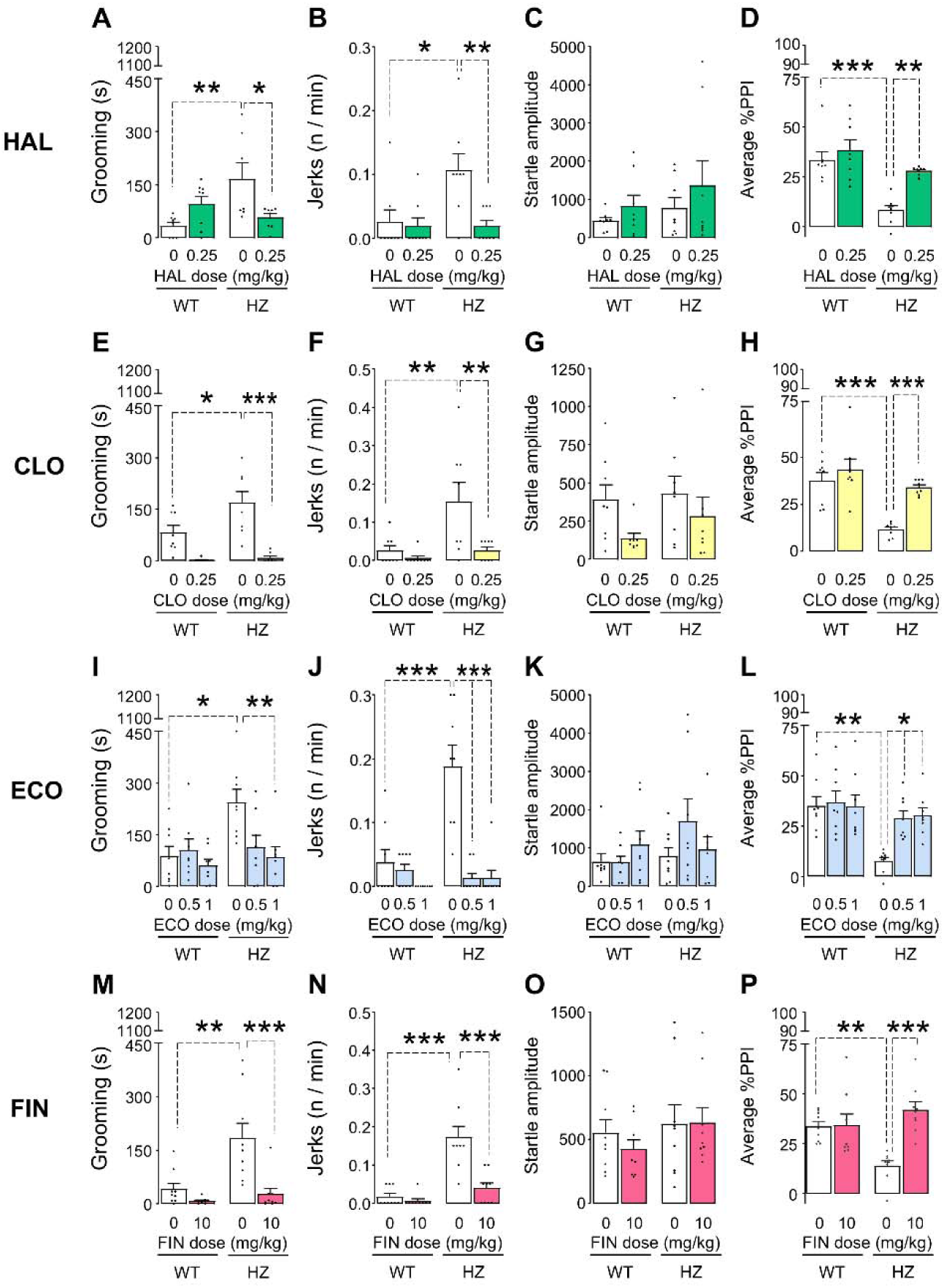
Tourette syndrome therapies are effective against the spontaneous tic-like behaviors of *Celsr3* heterozygous (HZ) mice. (A-D) The antipsychotic haloperidol (HAL; 0.25 mg/kg, IP, injected 45 min before the behavioral assessment) rescued the spontaneous increases of grooming duration (A) and jerk frequency (B), as well as the PPI deficits, without affecting the startle reflex amplitude (C-D). (E-H) The α2 agonist clonidine (CLO; 0.25 mg/kg, IP, injected 30 min before testing) elicited similar effects, as shown by the analyses of grooming duration (E), jerk frequency (F), startle amplitude (G), and PPI (H). (I-L) Similar effects were induced by the D_1_ receptor antagonist ecopipam (ECO; 0.5-1.0 mg/kg, IP, injected 20 min before testing), which reversed the spontaneous alterations of *Celsr3* HZ mice with respect to grooming duration (I), frequency of jerks (J), and PPI without affecting startle responses (K-L). (M-P) The 5α-reductase inhibitor finasteride (FIN; 10 mg/kg, IP, 30 min prior to testing) reduced the spontaneous grooming (M) and the frequency of jerks (N), while reversing the PPI deficits in *Celsr3* HZ mice without affecting acoustic startle amplitude (O-P). All experiments were performed in male animals and were analyzed by two-way ANOVA, with genotype and treatment as factors, followed by post-hoc analyses with Tukeýs test (*, P<0.05; **, P<0.01; ***, P<0.001). All data are shown as means ± SEM (n=8-9/group). For further details, see text and Suppl. Results.

### TS-related deficits in *Celsr3* HZ male mice are exacerbated by stress and reduced by benchmark TS therapies

To evaluate the predictive validity of adult *Celsr3* HZ mice as a TS model, we tested whether relevant behaviors could be modulated by environmental or pharmacological interventions known to affect tic severity in TS patients. First, we examined the effects of spatial confinement, a known stressor that exacerbates tic-like behaviors in TS models^41, 42^. Under this condition, both juvenile male and female HZ mice showed increased grooming, with males displaying a more pronounced response (Suppl. Fig.4A). Male HZ mice also exhibited a high frequency of body jerks regardless of stress (Suppl. Fig.4B). While startle amplitude increased across all males, only unstressed *Celsr3* HZ males showed PPI deficits, potentially due to a floor effect from stress in all groups (Suppl. Fig.4C-D).

Next, we assessed the effects of drugs known to reduce tics in TS. The D_2_ receptor antagonist haloperidol (0.25 mg/kg, IP) normalized grooming, head jerks, and PPI without affecting startle amplitude (Figs.2A-D). Similar improvements were observed with the α_2_ receptor agonist clonidine (0.25 mg/kg, IP; Figs.2E-H), the D_1_ receptor antagonist ecopipam (0.5-1.0 mg/kg, IP; Figs.2I-L), which is known to reduce tics in TS patients^43^, and the 5α-reductase inhibitor finasteride (10 mg/kg, IP; Figs.2M-P), effective in both patients and other tic disorder models^42,44,45^. Taken together, these findings suggest that most TS-like manifestations in *Celsr3* mutants were exacerbated by acute stress and mitigated by several therapies for TS patients.

Bulk transcriptomic studies identified significant changes in RNA processing and cytoskeleton regulation in striatal tissues from *Celsr3* HZ mice. Building on these findings, we directed our analyses toward molecular investigations of the dorsal striatum. Immunostaining confirmed that CELSR3 protein is expressed in this brain region, albeit at lower levels than the cortex, with HZ mice exhibiting a significant reduction in its levels (Suppl. Fig.5). We then examined the transcriptomic differences in bulk striatal tissue between WT and HZ littermates (n=8/genotype) (Suppl. Figs.6A-B). Unbiased clustering of differentially expressed genes (DEGs) distinctly differentiated between WT and HZ (Suppl. Fig.6C). As expected, *Celsr3* was one of the top genes whose expression was reduced in HZ mice. Other significantly altered genes were implicated in Wnt signaling and cell cycle progression, including *Tmed2*, *Chaserr*, *Slc7a5*, *Rpph1*, and *Ap4s1* (Suppl. Table 1). Gene ontology (GO) analysis found “Cytoskeleton’s Structure”, “Molecular Structure”, and “Cell-to-Cell Connections” as the most downregulated categories. In contrast, "mRNA Processing", "RNA Splicing", and "RNA Splicing Through Transesterification Reactions" were the most upregulated pathways (Suppl. Fig.6D and Suppl. Table 2).

### Spatial transcriptomic analyses of the striatum revealed changes in the extracellular matrix and inflammatory pathways

To further investigate the transcriptional variances across distinct territories of the striatum, comprehensive spatial transcriptomic analyses were conducted using 50-μm spots in coronal brain slices sectioned from brains of WT and *Celsr3* HZ mice (n=2/genotype) between -0.5 and +0.5 mm relative to bregma (anteroposterior) (Suppl. Fig.7). Seven spatial clusters, all enriched in *Drd1* and *Drd2* expression, were identified within the striatum and classified based on anatomical criteria and spatiomolecular markers^46^ (Figs.3A-C): 1) the dorsolateral striatum, characterized by high expression of *Nefm* and *Cnr1*; 2) the dorsomedial striatum, enriched in *Crym* and *Col6a1*; 3) the central striatum, highly enriched in markers related to white matter, such as *Mobp* and *Mag*; 4) the ventral striatum, showing enrichment for *Wfs1* and *Syt10*; 5) a cluster made up of intermittent punctate regions with high enrichment in interneurons positive for somatostatin, NPY, and nNOS (SNNINs); 6) intermittent puncta enriched in markers of striatal cholinergic interneurons (CINs), such as *Chat* and *Slc17a8*; and 7) a cluster of puncta enriched in markers of perivascular cells (*Vwf*, *Igf2*), such as endothelial cells and pericytes (Suppl. Table 3). We then analyzed the main GO pathways affected by genotype differences across all seven striatal territories. Pathway analysis revealed marked upregulation of several pathways related to chemotaxis of immune cells and inflammation in the dorsolateral striatum and SNNIN-enriched puncta, including “Cilium Movement,” “T Cell Chemotaxis,” “Neutrophil Chemotaxis,” “Granulocyte Chemotaxis,” and “Inflammatory Response” (Fig. 3D) are highlighted, along with a global downregulation of "Collagen-Containing Extracellular Matrix." which was prominently observed in both dorsolateral and ventral striatum (Fig.3E). A detailed analysis of the transcriptomic alterations observed in every single striatal cluster is presented in Supplemental Results (see also Suppl. Tables 4-6 for overall comparisons, upregulations, and downregulation in *Celsr3* HZ mutants).

**Fig. 3.**
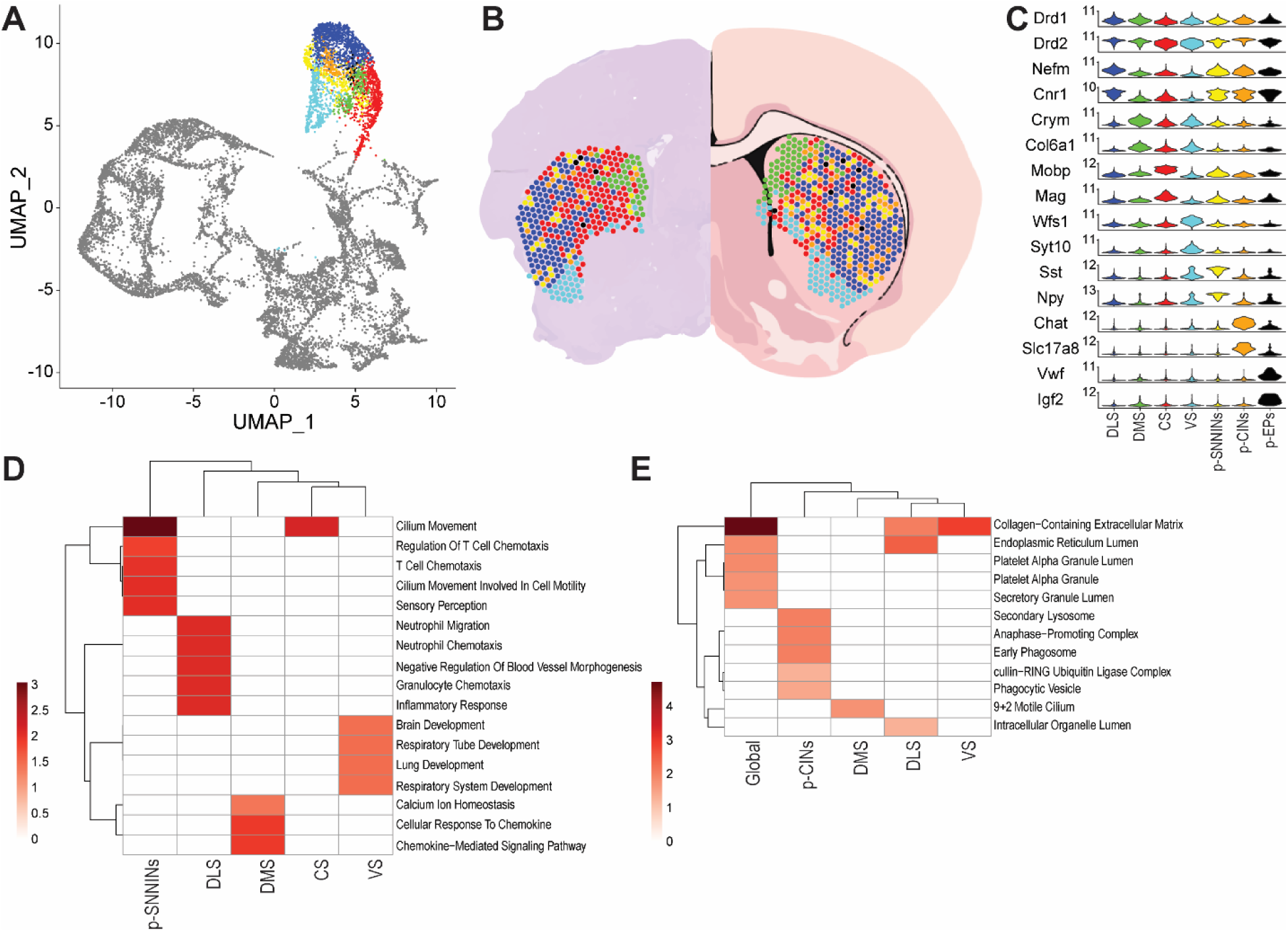
Spatial Transcriptomic Analyses of the Striatum in *Celsr3* heterozygous (HZ) mice reveal inflammatory alterations and reductions in collagen-containing matrix. (A) Uniform Manifold Approximation and Projection (UMAP) clustering of spatial transcriptomic data, showing seven distinct spatial clusters corresponding to the striatum (represented in color). (B) Spatial mapping of the seven clusters onto the mouse striatal section. The left side of the panel displays the placement of the dots on an actual brain slice, while the right panel shows the dots overlaid on a schematic representation of brain regions, illustrating the anatomical distribution of the distinct striatal clusters. (C): Violin plots of selected genetic spatiomolecular markers, demonstrating their differential expression across the seven identified clusters: dorsolateral striatum (DLS); dorsomedial striatum (DMS); central striatum (CS, enriched in white matter); ventral striatum (VS); puncta enriched in somatostatin-NPY-NOS1 positive interneurons (p-SNNINs); puncta enriched in cholinergic interneurons (p-CINs); and puncta enriched in perivascular cells (p-EPs). (D) Heatmap showing the upregulation of several Gene Ontology (GO) Biological Process pathways in striatal clusters of *Celsr3* HZ mice. (E) Heatmap representing the global downregulation of GO Cellular Component pathways in striatal clusters of *Celsr3* HZ mice. For further details, see text and Suppl. Results.

Single-nucleus transcriptomic studies highlighted subtle changes in striatal projection neurons (SPNs), interneurons, and microglia of *Celsr3* HZ mice. To identify which cell-specific aberrances are related to partial *Celsr3* deficiency, we next conducted single-nucleus-transcriptomic analyses of striatal tissues (n=3/genotype). Using established cell markers, we categorized seven cell types: SPNs, interneurons, oligodendrocytes, astrocytes, microglia, ependymal, and endothelial cells (Suppl. Figs.8-9 and Suppl. Tables 7-8). SPNs, interneurons, and microglia displayed clear gene expression differences, while oligodendrocytes and other cell types showed minimal transcriptomic variation, suggesting lesser impact from *Celsr3* deficiency (Suppl. Fig.8C).

GO analysis revealed changes in pathways like “Negative Regulation of Protein Phosphorylation” in SPNs, “Axon Extension,” “Cell-Cell Adhesion,” and “Neuron Projection” in interneurons, and “Synapse Pruning” and “Positive Regulation of Chemotaxis” in microglia (see Suppl. Table 8 and Suppl. Fig.10). To pinpoint affected neuronal subpopulations, we used Gamma-Poisson modeling on our snRNA-seq data (Suppl. Figs.11-13), identifying five SPN and five interneuron clusters (Figs.4A-B and Suppl. Tables 9-12). SPN clusters included D_1_-positive and D_2_-positive striosomal and matrisomal SPNs and a distinct eccentric “exopatch” (e-SPN) cluster, characterized by genes like *Col11a1*, *Tshz1*, *Otof*, and *Casz1*, which aligns with D_1_-positive striosomal neurons. At an adjusted p-value threshold of < 0.05, the predominant global signals within the GO Biological Process category indicated significant upregulation in pathways related to neuron projection development (Fig.4C and Suppl. Table 11), reflecting enhancements across most SPN families and SNNINs. Additionally, pathways associated with "Protein Localization to Microtubules" were upregulated specifically in SNINNs, parvalbumin-positive GABAergic interneurons (PVINs), and e-SPNs. Consistent with these observations, GO Cellular Component differences revealed differences in SPN clusters on "Neuron Projection," "Axon," and "Dendrite" pathways (Fig.4D and Suppl. Table 11). In contrast, calretinin-positive GABAergic interneurons (CRINs) exhibited selective downregulations in "Dendrite" and "Neuron Projection" pathways (Suppl. Fig.13). Lastly, GO Molecular Function analysis highlighted global regulation across several pathways, such as "Phosphatidylinositol Binding" and "Tubulin Binding," along with marked upregulation in pathways associated with ion channel functions in PVINs and "Adenylate Cyclase Regulation" in both PVINs and sD1-SPNs, suggesting increased activity in these cells (Fig.4E and Suppl. Table 11).

**Fig. 4.**
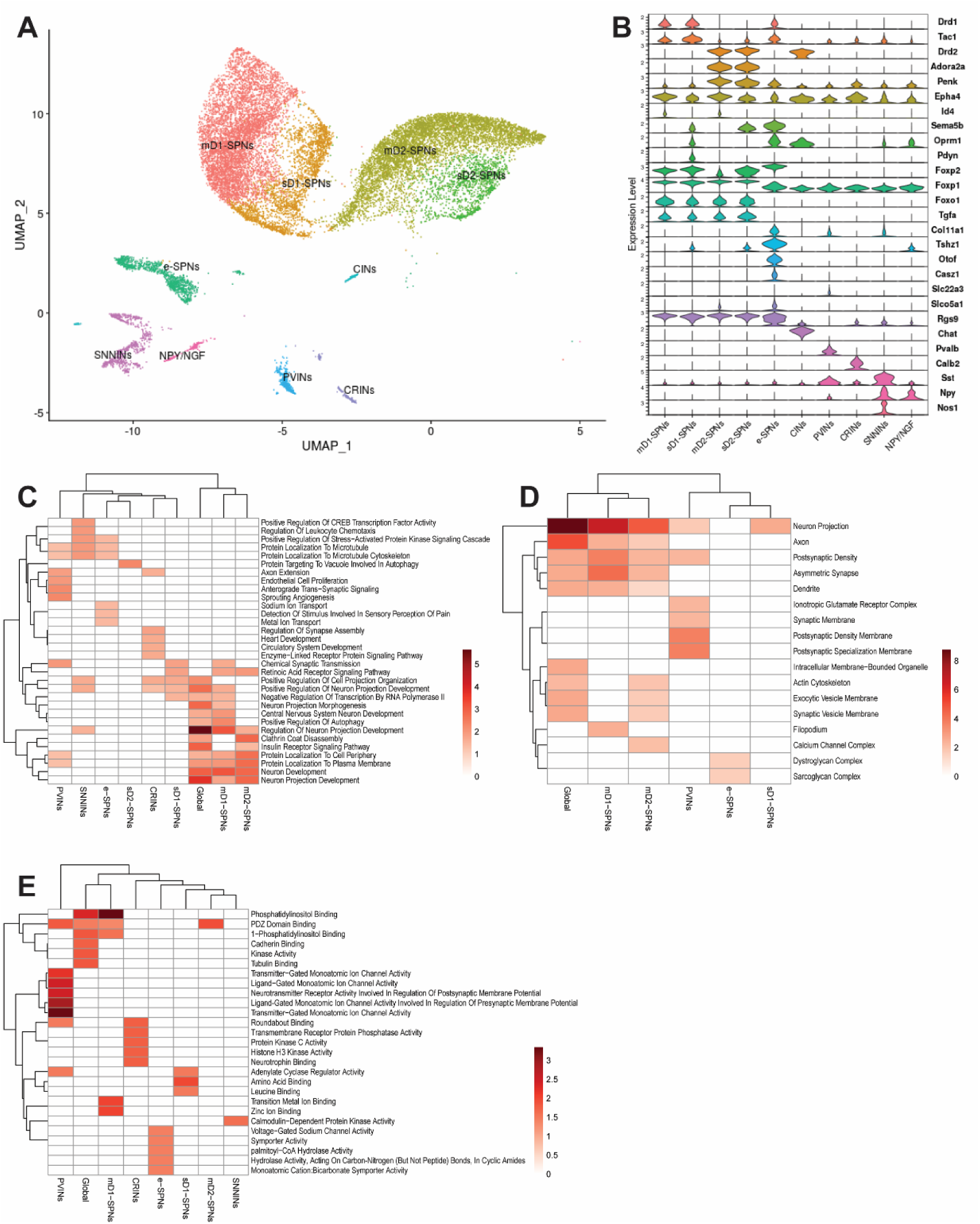
Single-nucleus transcriptomic analyses of neuron-associated clusters of *Celsr3* heterozygous (HZ) and wild-type (WT) reveal alteration in microtubule organization and signaling in select neuronal clusters. (A) Uniform Manifold Approximation and Projection (UMAP) of projection neurons and interneuron populations following SCTransformation with the generalized linear model, Gamma Poisson method. The resolution of the clusters was chosen when it resolved into distinct clusters and evaluating clusters in clustree. (B) Violin plots of marker genes used to distinguish the select striatal clusters differentiating known cell types. (C-E) Gene ontology analyses highlighting the most upregulated pathways among these cell clusters (C – GO Biological Process; D – GO Cellular Component; E-GO Molecular Function). Abbreviations: mD1-SPNs, matrisome-associated, D_1_-positive striatal projection neurons; sD1-SPNs, striosome-associated, D_1_-positive striatal projection neurons; mD2-SPNs, matrisome-associated, D_2_-positive striatal projection neurons; sD2-SPNs, striosome-associated, D_2_-positive striatal projection neurons; e-SPNs, eccentric striatal projection neurons (exopatch neurons); CINs, cholinergic interneurons; SNNINs, somatostatin-NPY-NOS1-positive interneurons; NPY/NGF, NPY neurogliaform interneurons; PVINs, parvalbumin-positive interneurons; CRINs, calretinin-positive interneurons. For further details, see text and Suppl. Tables and Results.

### Modulation of D_3_ dopamine receptors modifies grooming and tic-like behaviors in *Celsr3* HZ mice

Building on previous findings, we analyzed the top 20 transcripts within each neuron cluster, identifying a set of 35 differentially expressed genes (DEGs), including serine-threonine kinases, neurotrophin signaling molecules, myelin factors, and thyroid hormone-binding targets (Fig.5A). Notably, *Drd3* (encoding the D_3_ dopamine receptor) exhibited significant upregulation in sD_1_-SPNs and a concurrent reduction in CRINs (Fig.5A). Given prior evidence of enhanced dopamine release in the striatum of *Celsr3* mutants ^36^, we further investigated D_3_ dopamine receptors in the nigrostriatal system. We employed dopamine transporter (DAT) as a marker for dopaminergic fibers and assessed D_3_ receptor distribution via immunofluorescence and RNAscope *in situ* hybridization. Our analysis confirmed an increase in striatal cells coexpressing postsynaptic D_1_ and D_3_ receptors, while there was a reduction in presynaptic D_3_ receptors within dopaminergic fibers, indicated by decreased D_3_ and DAT coexpression. Conversely, no changes were observed in D_1_ and DAT levels (Figs.5B-I). In the substantia nigra *pars compacta*, we noted reduced DAT levels in *Celsr3* HZ mice, although overall D_3_ receptor expression and D_3_ levels in DAT-positive neurons remained consistent across genotypes (Suppl. Fig.14). These findings suggest that *Celsr3* deficiency impacts D_3_ dopamine receptor distribution in the striatum, supporting the hypothesis of altered dopaminergic signaling in this model.

**Fig. 5.**
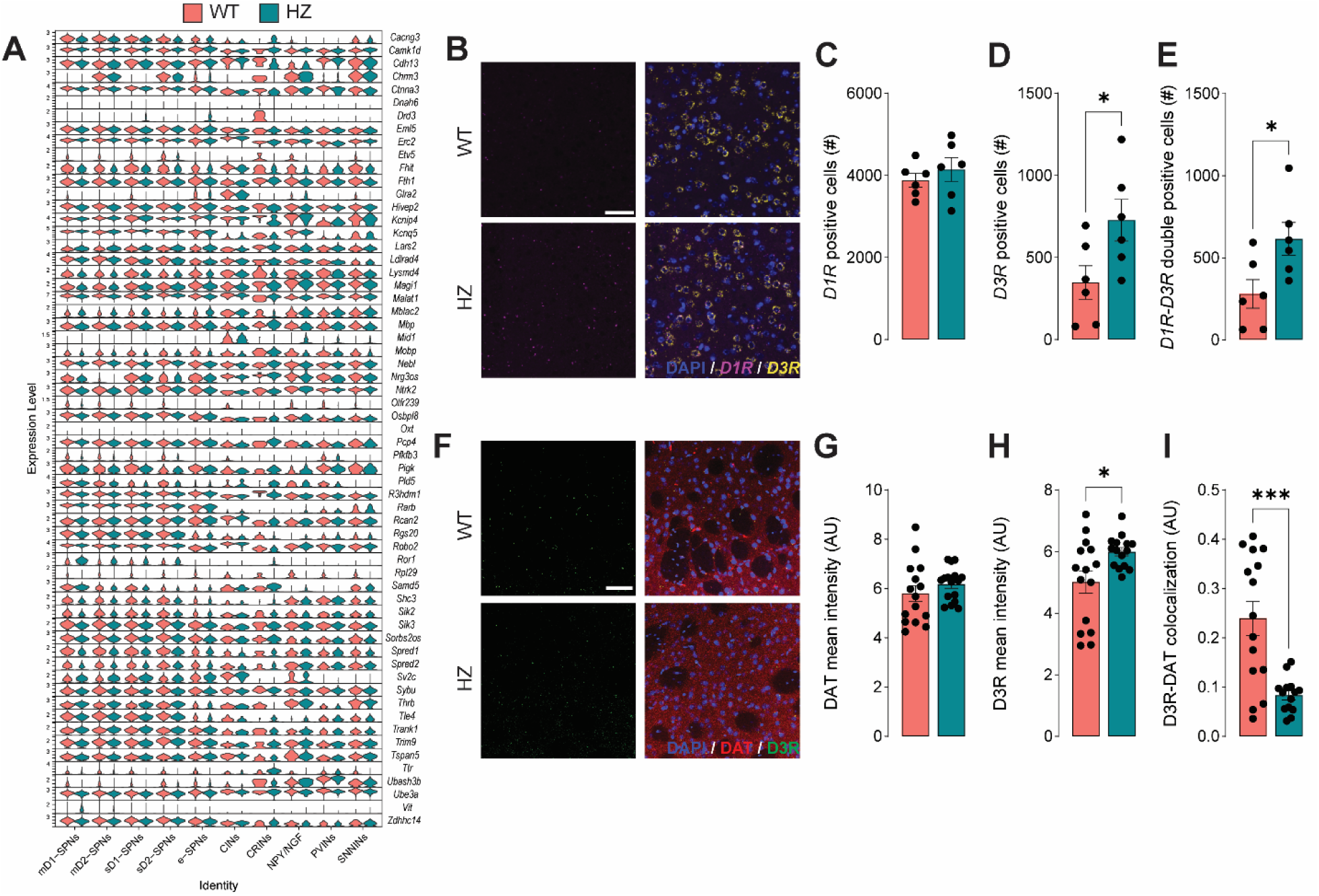
***Celsr3* heterozygous (HZ) mice exhibit alterations in striatal D_3_ dopamine receptors**. (A) Violin plots display the top differentially expressed genes (DEGs) across neuron clusters, highlighting genes expressed in at least two cell types. All striatal cell types are included. (B) Representative coronal sections from wild-type (WT) and HZ animals show *in situ* hybridization analysis of mRNA levels for D_1_ (yellow) and D_3_ (purple) receptors. (C-E) Quantification of *D1R* (C), *D3R* (D), and *D1R*-*D3R* double-positive cells (E). Semi-quantitative analyses using QuPath indicate an increase in *D3R*-positive and *D1R*-*D3R* double-positive cells. (F) Representative coronal sections from WT and *Celsr3* HZ animals showing immunofluorescent labeling of the D_3_ receptor (green) and dopamine transporter (DAT, red). (G-I) Semi-quantitative analysis of DAT (G), D3R (H), and DAT-D3R colocalization (I) was performed using ImageJ, revealing an increase in D3R mean intensity and a decrease in DAT-D3R colocalization. Merged images include DAPI (blue) for nuclear labeling. Scale bar: 50 µm. Data are presented as mean ± SEM (n=3/group). *, P<0.05; ***, P<0.001. For further details, see text and Suppl. Results.

In view of these findings, we hypothesized that alterations in postsynaptic D_3_ receptors in D_1_-positive neurons might contribute to the ontogeny of TS-related behavioral phenotypes in *Celsr3* HZ mice. To test this hypothesis, we measured the effects of the selective D_3_ receptor agonist PD-128907 (5 μg/kg, IP) and antagonist SB-277011A (10 mg/kg, IP) in TS-related alterations in these mutants. Activating the D_3_ receptors with PD-128907 increased grooming stereotypies in HZ, but not WT, mice (Fig.6A), dramatically increased tic-like responses in both genotypes (Fig.6B), while it did not significantly impact startle amplitude and PPI (Figs.6C-D). On the other hand, using the D_3_ receptor antagonist SB-277011A significantly reduced grooming in both WT and HZ mice (Fig.6E), reduced jerks in HZ, but not WT, mice (Fig.6F) without impacting startle amplitude or PPI (Figs.6G-H). Taken together, these data suggest that changes in cell-specific expression of D_3_ receptors in the dorsal striatum are implicated in the stereotypies and tic-like manifestations observed in HZ mice. In contrast, this receptor was not involved in the sensorimotor gating deficits exhibited by these mutants.

**Fig. 6.**
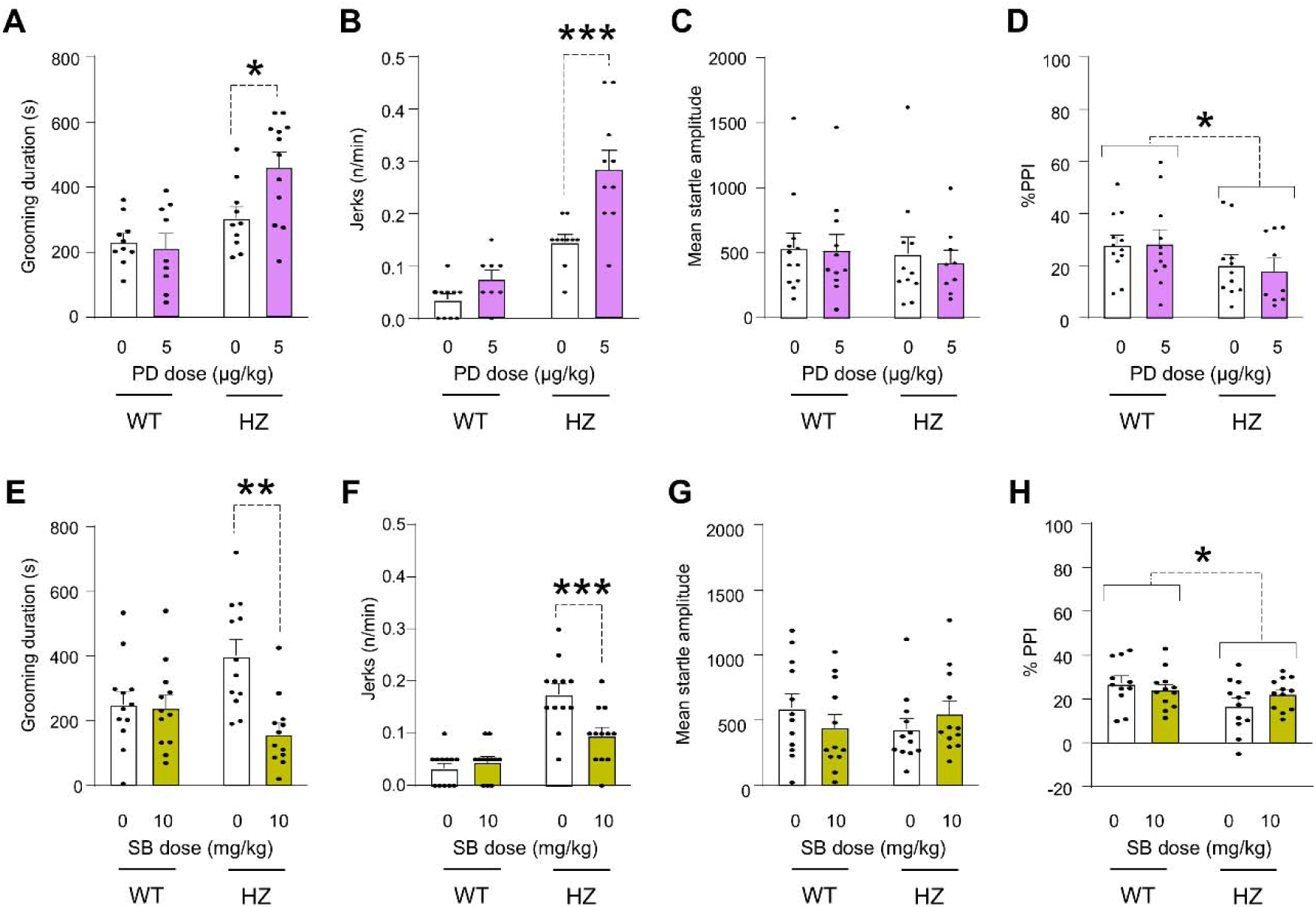
D_3_ dopamine receptor activation worsens tic-like behaviors in *Celsr3* heterozygous (HZ) mice, while antagonism reduces them. (A-H) Modulation of tic-related behaviors in *Celsr3* HZ mice treated either with the D_3_ receptor agonist PD-128907 (PD) or the D_3_ receptor antagonist SB-277011A (SB). All the alterations in behaviors spontaneously shown by *Celsr3* mice were exacerbated by PD treatment (A-D) and rescued by SB treatment (E-H). Data are shown as means ± SEM (n=9/group) and were analyzed by two-way ANOVA, with genotype and treatment as factors, followed by post-hoc analyses with Tukeýs correction (*, P<0.05; **, P<0.01; ***, P<0.001). WT, wild-type. For further details, see text and Suppl. Results.

## DISCUSSION

This study aimed to identify neurobehavioral and molecular signatures linked to *Celsr3* deficiency in animal models and assess their relevance to tic pathophysiology. In substantial agreement with recent findings on other mouse models harboring human-like *Celsr3* mutations ^36^, we found that the deficiency of this gene resulted in age- and sex-specific abnormalities resembling TS-related disturbances. While *Celsr3* HZ neonates behaved similarly to WT counterparts, weanling and juvenile mutants showed an increase in grooming stereotypies across both sexes, with this behavior becoming more pronounced in 42-day-old males; furthermore, juvenile male (but not female) HZ mice exhibited more axial jerks and PPI deficits. This sexual dimorphism in behavioral abnormalities appears to reflect the male predominance of TS and aligns with patterns seen in other TS mouse models^42, 48^. In contrast with our data, female mice with human-like *Celsr3* mutations (also maintained on a pure C57BL6/J background) were reported to exhibit greater grooming stereotypies than their male counterparts^36^. This discrepancy suggests that differences in sex predominance may stem from the specific *Celsr3* mutation, presumably in relation to its interaction with environmental factors.

Behavioral assessments in *Celsr3* HZ mice revealed elevated locomotor activity in both sexes and reduced object recognition in males. Although attentional deficits were not directly measured, these behavioral abnormalities align with those seen in rodent models of attention-deficit hyperactivity disorder (ADHD), a common comorbidity in TS patients. This suggests that *Celsr3* deficiency may contribute to behaviors relevant to both TS and other neurodevelopmental disorders. Further research is needed to clarify *Celsr3*’s role in attention and cognitive functions. Notably, *Celsr3* HZ mice did not show increased perseverative marble burying or anxiety-like behaviors, indicating that their stereotyped behaviors were not linked to abnormalities in these domains.

Consistent with evidence on the negative impact of acute stress in TS^49^ and tic disorder models, spatial confinement, a mild stressor known to increase corticosterone^41^ and brain allopregnanolone^50^, exacerbated certain behavioral abnormalities in *Celsr3* HZ mice. Notably, key TS-related behaviors were reduced by the TS-standard therapies haloperidol and clonidine^51^, as well as the D_1_ receptor antagonist ecopipam, recently shown to lower tic severity and improve global functioning in TS patients^43^. Additionally, finasteride, a steroidogenesis inhibitor known to reduce tic severity in open studies^44^, also reversed TS-related behaviors in *Celsr3* HZ mice. These drug effects were specific to *Celsr3* HZ mice, suggesting that alterations in catecholamines and neuroactive steroids may drive the TS-relevant behaviors in these mutants. Importantly, these effects were observed after acute administration, whereas therapeutic benefits in patients generally emerge with chronic treatment. This evidence further supports *Celsr3*-deficient animals as models for tic disorders with robust face, construct, and predictive validity. The presence of tic-like behaviors in *Celsr3*-deficient mice emphasizes the gene’s critical role in tic development, establishing these mutants as valuable models for studying TS pathophysiology and for testing potential new therapies.

To explore the molecular mechanisms whereby *Celsr3* deficiency leads to TS-relevant phenotypes, we analyzed the striatal transcriptomic profiles of HZ and WT mice through a combination of complementary approaches. Specifically, we conducted bulk, spatial, and single-nucleus transcriptomic analyses to examine gene expression changes across macro- and micro-(50-μm resolution) environmental milieus, as well as within specific cell types in the striatum. Collectively, these analyses pointed to an intricate spectrum of distinct yet complementary changes within the striatum. Bulk transcriptomics revealed a broad upregulation in biological processes linked to RNA processing and a significant downregulation in transcripts associated with microtubule regulation. Spatial transcriptomics identified microenvironmental changes across various striatal regions, highlighting molecular alterations in chemotaxis pathways and disruptions in collagen-containing extracellular matrix. Single-nucleus transcriptomic analyses further unveiled a complex array of transcriptomic signatures within several neuronal clusters and microglia, as well as alterations in the regulation of neuron projection development, as well as alterations in protein localization of microtubules and functional upregulation of sD1-SPNs and PVINs.

Overall, these findings suggest that *Celsr3* deficiency disrupts microtubule function, which plays a crucial role in both granulocyte chemotaxis ^52, 53^ and neurite growth ^29,30,54^. Although the direct impact of *Celsr3* on microtubules is not fully understood, the well-established role of this molecule in planar cell polarity (PCP) implies it may influence cytoskeletal dynamics. Supporting this idea, previous studies have shown that CELSR3 acts as a molecular guidepost, directing neuronal migration, axon guidance, and tract development ^55^.

Another key finding from our spatial transcriptomic analyses was the alteration of collagen-containing matrix across various striatal territories. This anomaly may reflect disrupted neutrophil function, as these cells play a well-documented role in degrading collagen and other extracellular matrix proteins^56^. Notably, several findings highlight collagen’s role in TS pathophysiology. For instance, the *COL27A1* gene, which encodes collagen type XXVII alpha 1, has been associated with TS susceptibility ^57, 58^. Collagen family proteins are essential for neuronal maturation, circuit formation, axon guidance, and synaptogenesis^59, 60^. Furthermore, deficiency in collagen VI (Col6) genes disrupts the dopaminergic system in mice^61^, underscoring collagen’s importance in the functional regulation of the striatum. It is also worth noting that the extracellular matrix influences microglial function^62^ and inflammation. Our single-nucleus transcriptomic analyses indicated that *Celsr3* deficiency impaired microglial capacity for synaptic pruning, a critical process for synaptic maturation^63^. Inflammation also significantly impacts motor control within corticostriatal circuits^64^ and is broadly implicated in TS pathogenesis^65, 66^.

Taken together, these results suggest that microtubular dysfunctions in *Celsr3*-deficient mice may contribute to several alterations across different striatal cell types. The resulting phenotypic changes are likely to contribute to aberrances in neuronal arborization and synaptogenesis, as well as inflammatory pathways involving neutrophils and microglia. The combination of these changes is likely to promote the formation of aberrant disinhibition foci in the dorsal striatum, ultimately increasing susceptibility to tics.

Among the multiple molecular changes observed in the striatum of *Celsr3* HZ mice, one of the most prominent was a striking imbalance in D_3_ dopamine receptor expression, with significant upregulation in sD_1_-SPNs and a parallel reduction in presynaptic D_3_ receptors in striatal dopaminergic fibers (as well as in striatal CRINs). Interestingly, no differences between genotypes were observed in D_3_ receptor distribution in the substantia nigra *pars compacta*, which hosts the cell bodies and proximal fibers of dopaminergic neurons projecting to the dorsal striatum. This suggests that *Celsr3* mutations likely impair D_3_ receptor translocation from somata to distal axonal projections in the striatum. The specific mechanisms underlying this altered D_3_ receptor distribution in the nigrostriatal pathway remain unclear; however, these changes may represent compensatory responses to broader disruptions in the dopaminergic system. Consistent with this possibility, DAT immunostaining indicated a pronounced reduction in dopaminergic fibers in the substantia nigra, echoing previous findings of disrupted dopaminergic fibers in *Celsr3* knockout embryos^32^. Notably, these fiber reductions were not evident in striatal presynaptic terminals, suggesting possible compensatory mechanisms within terminal fields that mitigate disruptions in dopaminergic fiber distribution. Together, the observed shifts in D_3_ receptor expression may significantly influence dopamine release dynamics, as D_3_ receptors—known for their high affinity for dopamine—play a crucial role in regulating dopaminergic signaling.

Irrespective of the processes underlying the imbalances in D_3_ receptor distribution, our studies revealed that D_3_ receptor activation intensified grooming and tic-like behaviors in *Celsr3* HZ mice, whereas D_3_ blockade alleviated these behaviors. These results, along with evidence that D_1_ and D_2_ receptor blockers also reduce tic-like behaviors, indicate a general hyperdopaminergic state in *Celsr3* HZ mice, consistent with prior findings of increased dopamine release in their striatum^36^.

In physiological conditions, D_3_ receptors are sparsely expressed in striatal neurons^67, 68^, while they are abundantly expressed in nigral dopaminergic neurons as well as their striatal projections, where presynaptic D_3_ receptors are posited to act as autoreceptors to limit dopamine release. In contrast with this scenario, our data suggest that, in *Celsr3* HZ mice, reduced presynaptic D_3_ receptor expression may drive excess dopamine release, while the upregulation of D_3_ receptors in striosomal D_1_-SPNs is expected to dysregulate dopamine signaling and trigger tic-like jerks and stereotypies. This idea aligns with evidence linking D_3_ receptor overexpression in striatal cells to dyskinetic manifestations ^69-71^, possibly through the formation of heteromeric receptors in combination with D_1_ receptors^72^. Further research is needed to clarify how D_3_ receptor upregulation in striosomal D_1_-SPNs contributes to tic-like behavior development.

Several limitations of our findings merit consideration. Firstly, we did not establish that specific activation of D_3_ receptor in sD_1_-SPNs is the proximal cause of tic-like behaviors in *Celsr3* HZ mice. A primary challenge in demonstrating this effect lies in the absence of an unequivocal marker for striosomal cells, as µ-opioid receptor (MOR), NR4A1, and other common patch markers are also distributed in the matrix, albeit at lower levels. Furthermore, the widespread presence of D_1_-positive SPNs throughout the striatum, in both soma and dendrites, reduces the precision of standard colocalization analyses with c-FOS or other activity markers for identifying specific activation of D_1_-SPNs. Future studies involving D_1_ reporter mice and chemogenetic constructs for the activation of specific subpopulations of SPNs are warranted to settle this critical issue and dissect the specific contribution of specific receptors and neurons in tic ontogeny.

Secondly, while our results indicate various functional changes in *Celsr3* HZ mice that may contribute to the development of tics, including alterations in D_3_ receptor expression, they do not provide a precise mechanistic explanation of how *Celsr3* alterations contribute to the pathogenesis of neurodevelopmental disorders. Future research endeavors should aim to uncover the specific striatal alterations and downstream mechanisms that underpin the modifications observed in *Celsr3*-deficient models. Understanding the precise role of *Celsr3* in TS may contribute to developing targeted therapeutic interventions for individuals with this complex neurodevelopmental disorder.

Despite these limitations, our study suggests that the predisposition to tic disorders associated with CELSR3 deficiency stems from a constellation of complex structural and functional changes within the nigrostriatal dopaminergic system and striatal organization. In this context, our findings also emphasize that combining deep behavioral phenotyping with multilevel transcriptomic analyses in animal models provides a powerful approach for uncovering the pathophysiological basis of tic-related responses driven by TS-associated mutations.

## MATERIALS AND METHODS

### Animals

*Celsr3* mutant mice in C57BL/6J background were generated as previously reported ^39^. As homozygous *Celsr3* knock-out mice die rapidly after birth^40^, all studies were performed using HZ and WT littermates, generated by breeding HZ sires with WT mothers (for genotyping details, see Supplemental Methods). Given that the identified *de novo CELSR3* mutations in humans typically affect only one allele^26-28^, the HZ genotype aligns with the alterations identified in TS-affected subjects. Housing facilities were maintained at 22°C, with lights on at 6:00 AM and lights off at 6:00 PM. Experimental manipulations were carried out between 9:00 AM and 4:00 PM. Every effort was made to minimize the number and suffering of animals. Thus, animal numbers for each experiment were determined through power analyses conducted on preliminary results. All experimental procedures were approved by the local Institutional Animal Care and Use Committees.

### Drugs

The following drugs were tested in WT and *Celsr3* HZ male mice: haloperidol, clonidine, ecopipam, finasteride, PD-128907, and SB-277011A. See Suppl. Methods for preparation and administration details.

### Behavioral assays

As shown in Suppl. Fig.1, behavioral assessments in *Celsr3* HZ mice and WT littermates were performed on males and females in three early developmental stages: newborn pups (postnatal days 1-7), weanlings (postnatal day 21), and juvenile (postnatal days 42-49) mice with different groups of mice to avoid developmental carryover effects. Behavioral assessment in newborn pups included righting and negative geotaxis reflexes, ultrasonic vocalizations (on postnatal day 6), and locomotor activity (on postnatal day 7)^73^.

In weanlings and juvenile mice, we evaluated spontaneous TS-relevant behavior, focusing on grooming, tic-like jerks, eye blinking, and prepulse inhibition of the acoustic startle. Testing was carried out as previously described^71^. In addition, juvenile mice were tested for behavioral responses related to TS-comorbid diagnoses. Paradigms included open-field locomotor activity, novel object recognition, marble burying, elevated plus maze, and wire-beam bridge (for more details, see Suppl. Methods). Each animal was randomly assigned to no more than three paradigms with at least 3-day intervals between tests to avoid carry-over effects.

Finally, separate cohorts of juvenile *Celsr3* HZ and WT male mice were used to study the impact of stress and the response to pharmacological treatments. To study the impact of stress, we subjected mice to spatial confinement, given that this mild stressor was shown to exacerbate TS-relevant responses in other mouse models^41, 42^. To ensure deep phenotyping of fine motoric abnormalities, each test was conducted with at least two separate cameras with different orientations, and videos were monitored by personnel blinded to genotypes and treatments with a 0.25-0.5 × speed. Each animal was assigned to only one treatment. All procedural details are presented in the Suppl. Methods.

### Bulk transcriptomic analyses

RNA was extracted from 16 male individuals (n=8/genotype). Sequencing and transcriptomic analyses were performed as previously described^74^ (see Suppl. Methods for details).

### Spatial Transcriptomic Preparation and Analysis

Male animals from each genotype underwent intracardial perfusion and brains were fixed to be processed for embedding. Spatial transcriptomics was performed on formalin-fixed, paraffin-embedded (FFPE) mouse brain sections. After RNA quality assessment and H&E staining and visualization, coronal slices (between -0.5 and +0.5 mm from bregma) were evaluated, prepared, and processed using the Visium Cytassist Spatial Gene Expression protocol (10X Genomics, Pleasanton, CA), with sequencing on a NovaSeq platform (Illumina, San Diego, CA). The data were aligned to the mm10 (GRCm38) genome and processed using Seurat (4.1.3) for quality control, filtering out low-quality spots. Dimensionality reduction and clustering were performed using SCTransformation and a Gamma-Poisson model, with a resolution of 1.2 to achieve detailed anatomical insights. The top 3000 spatially variable genes were identified for both WT and HZ samples, with pathway and Gene Ontology analyses performed via enrichR. Additional details are available in the Suppl. Methods.

### Single Nuclei Preparation and Transcriptomic Analyses

Freshly dissected striata were flash-frozen and processed for nuclei extraction in ice-cold PBS with inhibitors to protect RNA integrity. Tissues were homogenized, filtered, and prepared for library construction and sequencing (10X Genomics). Data were processed via Cell Ranger pipeline (10X Genomics), filtering out cells with low gene counts or high mitochondrial content, while excluding mitochondrial, hemoglobin, and bias genes. Dimensionality reduction and clustering were performed using Seurat (4.1.3), with a resolution of 0.4 to allow fine separation of interneuron clusters. Cell subpopulations were annotated by differential gene expression, validated with SingleR and enrichR, and analyzed for pathways and gene ontology using enrichR. For further methodological details, refer to the Suppl. Methods.

### *In situ* hybridization (ISH)

RNAscope™ ISH was performed on 5 µm FFPE coronal slices using an ACD RNAscope™ Multiplex Fluorescent V2 Assay (Advanced Cell Diagnostics, Inc., Newark, CA), with *MmDrd1* and *MmDrd3* probes in line with manufacturer’s instructions. Images were aquired on an Axioscan 7 Slide Scanner (Carl Zeiss Microscopy GmbH, Jena, Germany) and quantification was performed using QuPath software (0.5.1). For further methodological details, refer to the Suppl. Methods.

### Immunohistochemistry

For immunohistochemistry, 10 µm brain sections were deparaffinized, rehydrated, and staining was conducted with primary and secondary antibodies, with DAPI or hematoxylin counterstaining added for immunofluorescence and DAB-staining, respectively. Imaging was performed with a Leica SP8 confocal microscope (Leica Microsystem GmbH, Wetzlar, Germany) and AxioScan 7 Slide Scanner (Zeiss), with quantification via ImageJ (1.54i) and QuPath (0.5.1). Detailed protocols and antibody information are provided in the Suppl. Methods.

### Statistical analyses

For behavioral and biochemical data, normality and homoscedasticity of data distribution were verified using Kolmogorov-Smirnov and Bartlett’s tests. Statistical analyses were performed using one- or multiway ANOVA, as appropriate. Post-hoc comparisons were performed by Tukey’s test. Significance thresholds were set at 0.05. All methods for transcriptomic analysis and additional details are provided in Suppl. Information.

## Acknowledgments

We acknowledge the Huntsman Cancer Institute High-throughput Genomics and Bioinformatics Center, as well as the HSC Cell Imaging Core, for enabling sh-RNA sequencing and IF microscopy analyses. We thank Karen Odeh, Job Huxford, Elise Vandamme, and Easton Van Luik for their technical assistance with animal testing. We thank Anton Classen for his invaluable support in image acquisition and analysis.

## Data and Material Availability

All data needed to evaluate the conclusions in the paper are present in the paper and/or the Supplementary Materials. The data generated for sn-RNA seq are available on Gene Expression Omnibus (GSE249902 and GSE280709).

