## Supplementary Information for "Tic-related behaviors in *Celsr3* mutant mice are contributed by alterations of striatal D_3_ dopamine receptors"

**SUPPLEMENTAL INFORMATION**

**SUPPLEMENTAL MATERIALS AND METHODS**

**Genotyping of *Celsr3* heterozygous (HZ) mice**. After sperm cryo-recovery, the colony was maintained on a C57BL/6J background by crossing heterozygous (HZ) with wild-type (WT) mice, obtaining both genotypes in every litter. PCR genotyping was performed on postnatal day 21 using the following primers: CCCCCTGAACCTGAAACATAAAATG, ACTTCAGCACTGCACCCGACTTAC, and GCCCTGCGAGAACTACATGAAATG, as previously detailed ^1^.

**Phenotyping of *Celsr3* HZ mice**. *Neonates:* All the tests described below were performed as previously described ^2^ on *Celsr3* HZ and WT mice before genotyping. Thus, the number of animals is different among groups. From post-natal day (PND) 1 through postnatal day (PND)7, mice were tested for righting reflex. Briefly, pups were placed on their backs. and the latency to regain the natural position was recorded. On PND6, mice were tested for maternal separation-induced ultrasonic vocalizations (USVs). Recordings lasted 5 min, after which pups were immediately returned to the home cage and monitored for maternal reacceptance. Waveform Audio File Format (.wav) files were imported, and calls were automatically detected by *DeepSqueak* (v 2.6.2) within a frequency band of 20-120 kHz. On PND7, *Celsr3* HZ and WT mice were tested for locomotor activity, by placing them on a custom square open field for 5 min (12cm × 12cm), specifically designed for monitoring pup movements, and warmed at 32°C to prevent hypothermia and concomitant hypolocomotion in pups. Only 1 mouse/sex/litter was used in each group, to prevent potential litter effects.

*Weanlings*. *Celsr3* HZ and WT mice were tested immediately after weaning (PND21) to assess subtle alterations in spontaneous motor behaviors and deficits of sensorimotor gating, as previously described^3,4^. Behavioral tests on weanlings included the assessment of i) spontaneous stereotypies, including self-grooming, digging, and chewing; ii) tic-like axial jerks; iii) eye blinking activity; iv) prepulse inhibition (PPI) of the acoustic startle. To capture fine abnormalities, behavioral studies were conducted on 16-18 mice/group, and each spontaneous behavior was videorecorded from 2-4 distinct cameras placed in opposite directions and monitored by personnel (blind to genotype) at reduced speed (× 0.25 - 0.5). Only one mouse/sex/litter was used in each group, to prevent potential litter effects.

*Juvenile mice:* *Celsr3* HZ and WT mice were tested on PND42 to assess the same spontaneous TS-relevant responses reported above. Additionally, between PND43 and PND49, a comprehensive battery of behavioral tasks was used to capture manifestations akin to TS-related comorbid symptoms. Behavioral studies were conducted on 18 animals/group. Furthermore, given the role of acute stress in triggering TS-like manifestations, TS-like phenotypes were also evaluated in a separate cohort during/after spatial confinement, a mild stressor known to exacerbate tics and stereotyped responses in other animal models of TS ^3,4^. To produce this mild stressor, animals were confined within a clear, bottomless Plexiglas cylinder (10 cm in diameter × 30 cm in height), which was placed in a familiar cage, and deeply sunk into the bedding to ensure stability. Furthermore, to evaluate the face and predictivity of this model, the effects of different classes of treatments for tic disorders were also measured in male mice. The effects of acute stress and pharmacological studies were performed on 8-9 animals/group.

Before all behavioral testing, mice were consistently habituated to the experimental room and an empty adjacent room for at least 60 min per day. On the day of the test, mice were carried to the waiting room for at least 60 min, and then one animal at a time was brought to the experimental room using a clean transport cage. The waiting and testing rooms were kept at 100 lux (dim light). Behavioral tests performed on juvenile mice included the assessment of i) spontaneous and stress-induced stereotypies, including self-grooming, digging, and chewing; ii) tic-like axial jerks; iii) eye blinking activity; iv) PPI of the acoustic startle; v) marble-burying; vi) wire-beam elevated bridge; vii) novel object recognition; viii) open-field locomotor activity; and ix) elevated plus-maze. Testing was performed as described previously ^3,4^. Each spontaneous behavior was videorecorded from 2-4 distinct cameras placed in opposite directions and monitored by blinded personnel at reduced speed (× 0.25- 0.5). Only 1 mouse/sex/litter was used in each group, to prevent potential litter effects. Each animal was not subjected to more than three tests to avoid stress carry-over effects.

**Spontaneous stereotypies.** Self-grooming behavior and jerks of freely moving mice were assessed in a regular mouse cage (29 × 17 × 13 cm) with bedding material; instead, for the evaluation of the effect of a mild stressor, animals were confined within a clear, bottomless Plexiglas cylinder (10 cm in diameter × 30 cm in height). After a 10-min familiarization period to avoid neophobia-induced alterations, the behavior was video-recorded for 20 min. Grooming behavior (including complete and incomplete sequences of licking, scratching, and washing the paws, head, body, and tail) and number of jerks were scored by trained observers blinded to genotype and sex.

**Assessment of eye blinks and fine head movements.** Eye blinks were monitored during spatial confinement (as described above). The cylinder was mounted on a square platform adjacent to four video cameras placed on each side ^5^. This configuration allows remote and continuous monitoring of eye blinks in a non-invasive fashion and without the employment of head restraint bars. Animals were exposed to the apparatus for two days before testing. Eyeblinks were scored by trained observers blind to genotypes and other conditions.

**PPI of the startle reflex**. Startle testing was conducted in sound-attenuating ventilated startle chambers (SR-LAB, San Diego Instruments, San Diego, CA) as previously detailed ^3,4^. The acoustic PPI protocol featured a 70 dB background broadband noise (5 min acclimation period), followed by three consecutive blocks of "pulse", "prepulse + pulse", and "no stimulus" trials. During the first and third blocks, mice received five "pulse alone" trials of 115 dB. During the second block, animals underwent a pseudo-random sequence of fifty trials, consisting of "pulse-alone" trials (twelve events), pulses preceded by 73 (PP3), 76 (PP6), or 82 (PP12) dB "prepulses" (ten events for each level of prepulse), and "no stimulus" trials (eight events, consisting of background white noise delivery). Intertrial intervals (ITI) in both protocols were selected randomly between 10 and 15 s. A dynamic calibration system was used to ensure comparable sensitivities across chambers. Percent PPI (%PPI) was calculated using the formula (Mean Startle Amplitude, MSA, calculated as the mean peak voltage of the startle response to pulse-alone trials):

100 – *MSA* × “*pre-pulse pulse*” trials / *MSA* × “*pulse alone*” trials × 100

Since our initial characterization of *Celsr3* HZ mice showed a robust main effect for prepulse loudness levels (see Results) but no interactions between this factor and others, their values were averaged across the three prepulse loudness levels for all pharmacological studies. When PPI and startle reflex were measured in conjunction with confinement stress, animals were tested immediately after exposure to this stressor due to the incompatibility of startle chambers with confinement cylinders.

**Locomotor activity in the open field.** Horizontal locomotor activity was tested in a black Plexiglass open field arena (40 × 40 × 40 cm), as previously described^4^. Briefly, mice were placed in the center of the arena and allowed to explore freely for 5 min. Locomotor activity and time in the center (defined as a central 28.3 × 28.3 cm square) were scored and analyzed using behavioral tracking software (EthoVision XT, Noldus, Wageningen, The Netherlands).

**Novel object recognition test.** Testing was executed as previously indicated^6^. Mice were individually acclimatized to Makrolon cages for 15 min each. The day after, animals were exposed to two novel black plastic cylinders (8 cm tall × 3.5 cm in diameter), affixed to the floor, and symmetrically placed at 6 cm from the two nearest walls. Mice were placed in a corner, facing the center and at equal distance from the two objects. Their start position was rotated and counterbalanced for each genotype and sex throughout the test. 24 h later, mice were placed in the same cage. One of the cylinders was replaced by a novel plastic rectangular block (3 × 3 × 6 cm), which was placed in a counterbalanced fashion to avoid experimental bias. The behaviors for both sessions were videotaped for 15 min. Analysis included the number and total duration of exploratory approaches between novel and familiar objects. Exploration was defined as sniffing or touching either of the two objects with the snout; sitting on the object was not considered exploration. In the second exploration trial, an object exploration index was calculated as the ratio of the duration of the exploratory approaches targeting the novel object over the time of exploration of both objects.

**Marble-burying test.** Testing was executed as previously indicated^7^. Briefly, each mouse was transferred individually into a clean home cage (29 × 17 × 13 cm) containing 16 colored marbles homogenously distributed over 5 cm bedding for a 15-min trial. Testing was performed with normal illumination conditions (300 lux) for 15 min. Buried marbles were counted by blinded experimenters at the end of testing.

**Elevated plus-maze.** Anxiety-like behavior was assessed using an elevated plus-maze under conditions of normal environmental illumination (300 lux) as previously described^4^. The maze consisted of two open arms (35 × 6 cm) and two closed arms (35 × 6 × 21 cm) extending from a central platform (6 × 6 cm) elevated at 74 cm from the ground. Mice were placed in the center area of the maze with their heads directed toward a closed arm. The time spent in the open arms (with all four paws in the arm) was measured and expressed as a percentage of the total time in the maze. The number of entries and head dips were also monitored. The overall duration of the test was 5 min.

**Wire-beam elevated bridge test.** Testing was executed as previously indicated^7^. The apparatus consisted of a 50 cm high platform and a wire beam bridge 50 cm above ground that connected the platform to the end of the apparatus. The platform was 1 cm in length × 1 cm in width, surrounded by a 30 cm high wall on one side open facing the bridge. The bridge was 40 cm in length × 1 cm in width and consisted of two parallel beams (1 cm in diameter) perpendicularly connected by 40 equally distanced crossties. A foam mat was placed on the floor below the bridge to protect against the infrequent possibility that an animal fell off the bridge. Mice were individually placed on the starting point, and the latency to cross the bridge was measured using a 5-min cutoff time.

**RNA extraction and RNA-seq***.* RNA was extracted from striata of 16 male animals (8 mice/genotype) using the RNeasy Lipid Tissue Mini Kit (Qiagen, Hilden, Germany), and eluted in 40 µl of RNase-free water. RNA's quality control involved RIN value assessment via a Qubit (Invitrogen, Carlsbad, CA) and the 260/280 ratio using a Nanodrop (ThermoFisher Scientific, Waltham, MA). RNA extracted was sequenced using NovaSeq Reagent Kit v1.5_150x150 bp (100 M read-pairs) Sequencing (Illumina Inc, San Diego CA) and sequencing libraries were prepared using Illumina TruSeq Stranded Total RNA Library Prep Ribo-Zero Gold kit (Illumina).

**Bulk transcriptomic data processing**. Raw reads were aligned using *STAR v2.7.5b*^8^ using the reference genome GRCm39 and the count table was generated using the function *featureCounts*, as implemented in the R-package *subread*^9^. Read mapping was assessed using *MultiQC v1.12*^10^, and samples with less than 70% of uniquely mapped reads were removed before conducting downstream analysis. Raw counts were imported on *DESeq2*^11^. After removing genes with less than ten total counts, data were transformed with the variance stabilizing method^12^ before running principal components analysis (PCA). Outliers were defined as samples with PC1 or PC2 values above plus or minus three standard deviations from the mean in the first two rounds of PCA. The relationship between RIN and gene expression was conducted by correlating the top two principal components using Pearson's *r.* Raw counts were normalized using *DESeq2*, and differential expression analysis was performed between HZ and WT, including RIN as a covariate. P values were adjusted for multiple testing using the False Discovery Rate (FDR) method^13^. Genes with adjusted p < 0.05 were considered statistically significant differentially expressed genes (DEGs). Pathway analysis was conducted on the DEGs using *clusterProfiler* R-package^14^, referencing to the Gene Ontology (GO) database, and adjusting the p-values with the FDR method. Cell-specific expression enrichment was computed across the DEGs using the cell-specific gene markers obtained according to the previously described workflow^15^ using published single-cell RNA sequencing murine data^16^. The enrichment test was conducted by hypergeometric statistics using the function *enrichment* from the R-package *bc3net*^17^ and adjusting the P-values with the FDR method.

**Tissue collection.** Male animals from both genotypes were anesthetized with ketamine (80 mg/kg) and xylazine (15 mg/kg), followed by intracardial perfusion with phosphate-buffered saline (PBS) for 5 min. Brains were harvested and fixed in 10% formalin for 48 hours. Subsequently, the tissue was washed in 30% sucrose in PBS and then transferred to 70% ethanol prior to the embedding process. Formalin-fixed, paraffin-embedded (FFPE) tissues were sectioned at 5 µm using a microtome for spatial transcriptomics and *in situ* hybridization analyses, and at 10 µm for immunohistochemistry analyses.

**Spatial transcriptomic preparation and analysis.** Formalin-fixed paraffin-embedded (FFPE) mouse brains were sectioned and RNA quality assessed through DV200 measurements on an Agilent 2200 Tapestation (Agilent Technologies, Inc., Santa Clara, CA). For samples showing a DV200 ≥ 30%, additional sections were prepared, stained with H&E, and visualized using an Axioscan microscope (Carl Zeiss Microscopy GmbH, Jena, Germany) with a 20x objective under glycerol-stabilized coverslips. Multiple coronal sections for each animal, from -0.5 to +0.5 mm (AP) relative to the bregma, were examined. Optimal slides proceeded to spatial transcriptomics using the Visium Cytassist Spatial Gene Expression for FFPE protocol (10X Genomics Inc., Pleasanton, CA), followed by sequencing with the NovaSeq Reagent kit v1.5 50x50 bp (Illumina). The Space Ranger pipeline (10x Genomics) processed sequencing files and aligned them to mm10, M_musculus_Dec_2011, GRCm38.

Four samples, including two independent Celsr3 WT and HZ samples, were analyzed. Quality control and downstream analysis were conducted using Seurat (4.1.3) ^18–22^. Spots were filtered to exclude those with fewer than 500 genes, over 25% mitochondrial genes, or over 20% hemoglobin content. Dimensionality reduction and clustering employed SCTransformation (0.3.5) with the Gamma-Poisson generalized linear model (glmGamPoi, 1.4.0) ^23^. FindIntegrationAnchors^20^ was utilized to adjust for batch effects between the two animal samples for each group. PCA, FindNeighbor, UMAP, and FindClusters analyses were performed on the SCT “integrated” dataset, with a resolution of 1.2 to reveal anatomical detail in the striatal regions. This spatial analysis, based on 50-μm spots, occasionally precluded identification of individual cell subpopulations within clusters; however, differential gene expression highlighted spatial features at this resolution.

**Single nuclei preparation and transcriptomic analyses.** Freshly dissected striatal tissue was immediately flash-frozen in liquid nitrogen, embedded in Optimal Cutting Temperature (OCT; Tissue Plus, #4585, Fisher HealthCare, Waltham, MA) compound, and stored at -80°C prior to nuclei extraction. At the start of processing, samples were gently thawed in 2 ml of ice-cold phosphate-buffered saline (PBS; devoid of Ca²⁺ and Mg²⁺), to which the following supplements were added: 0.54 μM Necrostatin (Inc. #J65341, ThermoFisher Scientific Chemicals, Waltham, MA), 1 μM HPN-07 (SML2163, Sigma-Aldrich), 0.32 μM sodium hydroxybutyrate (A1161314, ThermoFisher Scientific Chemicals), 78 nM Q-VD-Oph (SML0063, Sigma-Aldrich, St. Louis, MO), and 0.2 U/ml of Ribolock RNase inhibitor (EO0381, Life Technologies Corporation, Carlsbad, CA). Tissue fragments, placed on dry ice in a 60-mm Petri dish, were minced to sizes under 2 mm with a razor blade. The tissue was then transferred to a 2 ml Dounce homogenizer containing 1.5 ml nuclei extraction buffer (130-128-024, Miltenyi Biotec, Bergisch Gladbach, Germany) with 0.2 U/ml Ribolock RNase inhibitor. The homogenization procedure involved ten strokes using pestle A, followed by ten strokes with pestle B, with a subsequent 5-minute incubation on ice. This was followed by twenty strokes with pestle B, a second 5-minute ice incubation, and a final twenty strokes with pestle B. Samples were centrifuged at 600×g for 5 minutes, resuspended in 400 μl PBS with 2% BSA, and clarified by sequentially passing through 70-μm and 40-μm FLOWMI strainers. Library construction and sequencing were subsequently performed using the 10X Genomics platform.

Data processing was carried out using the Cell Ranger Single Cell pipeline^18^ (10X Genomics), generating alignments and counts with default settings and the mm10 reference dataset for alignment. Downstream analysis and quality control were conducted using Seurat (4.1.3)^18–22^. Cells were filtered based on thresholds of less than 750 detected genes and mitochondrial gene expression greater than 15%. Genes were filtered out, specifically: mitochondrial genes, hemoglobin genes, and potential sources of bias, including Gm42418, AY036118, Gm47283, Rpl26, Gstp1, Rpl35a, Erh, Slc25a5, Pgk1, Eno1, Tubb2a, Emc4, Scg5, Ehd2, Espl1, Jarid1d, Pnpla4, Rps4y1, Xist, Tsix, Eif2s3y, Ddx3y, Uty, Kdm5d, Cmss1, AY036118, and Gm47283^24^.

Dimensionality reduction was achieved through nearest-neighbor identification and data clustering based on scaled RNA expression at a resolution of 0.1 to identify primary cell types. Subsequently, SCTransformation (0.3.5) with the Gamma-Poisson generalized linear model (glmGamPoi, 1.4.0)^23^ was applied, and a resolution of 0.4 was selected, as interneuron subpopulations began differentiating into distinct clusters. Differential expression analysis allowed for cell-type annotation of clusters, with manual annotations validated against automated classification using SingleR (1.6.1) and enrichR^25–27^ (3.1, CellMarker Augmented 2021). Pathway and Gene Ontology analyses were performed using enrichR (3.1) ^25–27^. Differential gene expression between Celsr3 HZ and WT control was assessed using FindMarkers. To identify genes clusters showing significant differential expression between Celsr3 HZ and WT controls in two or more clusters, genes within each cluster expressed in at least 10% of cells per cluster were analyzed. Poorly annotated genes (e.g., genes lacking canonical names beginning with "AC-", "Gm-", or "-Rik") were excluded.

**RNAscope™ *in situ* hybridization.** The ACD RNAscope™ Multiplex Fluorescent V2 Assay (#323100, Advanced Cell Diagnostics, Newark, CA) and probes (MmDrd1, #461901; MmDrd3, #447721-C2) were used according to manufacturer’s instructions. Briefly, 5 μm coronal sections were deparaffinized, rehydrated, treated with hydrogen peroxide to reduce background, put through a permeabilization/RNA retrieval procedure, and then sequentially, probes were hybridized, Opal dyes (Opal 620 and Opal 690, Akoya Biosciences, Marlborough, MA) conjugated to amplify the signal, washed and blocked, and finally coverslipped with DAPI Prolong Gold mounting medium (ThermoFisher Scientific). Images were acquired using Axioscan 7 Slide Scanner (Zeiss) using Opal optimized filters and quantified following the manufacturer's guidelines using *QuPath* (v 0.5.1)^28^, as outlined in the technical note, "Using QuPath to analyze RNAscope™, BaseScope™ and miRNAscope™ images," available at <https://acdbio.com/qupath-rna-ish-analysis>.

**Immunohistochemistry.** Brain slices (10 µm) were deparaffinized through three washes in xylene and rehydrated in a series of ethanol solutions with decreasing concentrations. Antigen retrieval was performed by incubating the sections in Tris-EDTA Antigen Retrieval Buffer (Proteintech, Rosemont, IL) at 95°C for 15 min, followed by washing in tris-buffered saline (TBS: 50 mM Tris-HCl, pH 7.4, 150 mM NaCl). For DAB staining, endogenous peroxidases were quenched using a solution of H_2_O_2_ and methanol; this step was omitted for immunofluorescence analysis. Slides were then incubated in TBS-A (TBS + 0.1% Triton-X) and subsequently in TBS-B (TBS-A + 2% BSA). Primary antibodies (details listed below) were diluted in TBS-B and incubated overnight at 4°C. The following day, excess antibodies were removed by washing the slides twice in TBS-A, followed by one wash in TBS-B. Secondary antibodies were diluted in TBS-B and incubated for one hour at room temperature. For immunofluorescence, this incubation was carried out in the dark, with DAPI counterstaining (Sigma-Aldrich) added to the secondary antibody (Goat anti-Rabbit IgG Alexa Fluor 555 and AlexaFluor 647, ThermoFisher Scientific). Slides were then washed in TBS and mounted with Fluoroshield mounting medium (Sigma-Aldrich). For DAB staining, following incubation with the secondary antibody (included in the kit VECTASTAIN® Elite® ABC-HRP Kit, Vector Laboratories, Newark, CA), slides were washed in TBS-A, followed by TBS-B, and incubated with the ABC Elite system (Vector Laboratories) for one hour at room temperature. Slides were then washed three times in TBS before being exposed to DAB solution (Vector Laboratories) for 4 min. The DAB reaction was halted with water, and after three washes in water, slides were counterstained with Harris hematoxylin (Polysciences, Inc., Warrington, PA), dehydrated, and mounted with DPX mounting medium (Sigma-Aldrich).

Immunofluorescence images were captured using a Leica SP8 confocal microscope (Leica Microsystems, Wetzlar, Germany), and quantifications were performed using ImageJ (1.54i) software ^29^. DAB-stained sections were imaged with an AxioScan 7 Slide Scanner (Carl Zeiss), and quantification was conducted using QuPath (0.5.1) ^28^.

**Antibodies.** The following antibodies were used in the presented study: Anti-Dopamine Receptor D3/DRD3 antibody (ab42114, Abcam, Cambridge, UK), DAT Polyclonal antibody (22524-1-AP, Proteintech), CELSR3 Polyclonal antibody (28835-1-AP, Proteintech).

**SUPPLEMENTAL RESULTS**

**Behavioral phenotyping of *Celsr3* HZ at different ages.** We first analyzed the behavioral responses in neonate *Celsr3* HZ mice, in contrast with WT counterparts. As shown in Suppl. Fig, 2, no major differences were found in righting reflex (Suppl. Fig. 2A). Similarly, ultrasonic vocalizations (Suppl. Fig. 2B) and locomotor activity (Suppl. Fig. 2C) were equivalent between genotypes at this developmental stage. We then conducted a battery of tests aimed at capturing fine behavioral abnormalities in weanlings (PND21). These analyses revealed that both male and female HZ mice exhibited greater levels of spontaneous self-grooming (Fig. 1A) [Main effect of genotype: F(1,68)=5.78; P=0.02]. Conversely, no differences in spontaneous body jerks (Fig. 1B), chewing movements (Fig. 1C), or eye blinks (Fig. 1D) were found. Female HZ mice engaged in significantly less digging behavior than their WT counterparts [Sex × Genotype interaction: F(1,68)=5.78; P=0.02; P<0.05 for post-hoc comparisons between WT and HZ females] (Fig. 1E). All groups displayed similar acoustic startle responses (Fig. 1F); in contrast, male, but not female, *Celsr3* HZ mice exhibited lower PPI values across all prepulse loudness levels than their WT littermates (Fig. 1G) [Sex × Genotype interaction: F(1,204)=5.78; P=0.02; P<0.01 for post-hoc comparisons between WT and HZ males].

At PND42, behavioral testing revealed a significant increase in self-grooming stereotypies in *Celsr3* HZ mice (Suppl. Fig. 1H) [Main effect of genotype: F(1,68)=10.46, *P*=0.002; sex × genotype interaction: F(1,68)=4.11, *P*=0.046; *P*<0.01 for comparison between male HZ and WT mice]. Similarly, *Celsr3* HZ males exhibited a higher frequency of head jerks (Suppl. Fig. 1I) [Genotype × Sex interaction: F(1,68)=4.06, *P*=0.048; *P*=0.005 and *P*<0.01 for post-hoc comparisons between male HZ vs male WT and vs female HZ, respectively]. No significant differences were observed in spontaneous chewing, eye blinking, or digging (Suppl. Figs 1J–1L). *Celsr3* HZ males, but not females, demonstrated notable PPI deficits, which were consistent across all prepulse levels (Suppl. Fig. 1N) and were accompanied by changes in acoustic startle amplitude (Suppl. Fig. 1M), with female mice showing a lower startle amplitude reflex compared to males, regardless of genotype [Main effect of sex: F(1,68)=8.86, *P*=0.004].

To further investigate the behavior of HZ mice, we assessed additional paradigms related to locomotor activity, cognition, anxiety, and compulsivity. *Celsr3* HZ mice exhibited increased locomotor activity regardless of sex, as indicated by the greater distance covered (Suppl. Fig. 3A) [Main effect of genotype: F(1,68)=4.38, *P*=0.04] and the increased time spent in the center of the arena (Suppl. Fig. 3B) [Main effect of genotype: F(1,68)=6.46, *P*=0.01]. Additionally, *Celsr3* HZ males, but not females, showed a significant reduction in novel object recognition (Suppl. Fig. 3C) [Sex × genotype interaction: F(1,60)=6.19, *P*=0.01; *P*<0.01 for comparisons between male HZ and WT mice]. No differences in marble-burying (Suppl. Fig. 3D), elevated-plus-maze behavior (Suppl. Figs. 3E–3G), or latency to cross a wire-beam bridge (Suppl. Fig. 3H) were observed, suggesting no overt alterations in compulsivity, environmental anxiety, or impulsivity.

After characterizing spontaneous behavior, we tested the effects of spatial confinement, a mild stressor known to exacerbate tic-like responses in other mouse models of Tourette Syndrome (TS) ^3,4^. Both male and female HZ mice displayed a significant increase in grooming responses, which was further heightened by stress (Suppl. Fig. 4A) [Main effect of genotype: F(1,64)=46.92, *P*<0.0001; main effect of stress: F(1,64)=19.85, *P*<0.0001]. This effect was more pronounced in males [Stress × sex interaction: F(1,64)=7.56, *P*=0.008; *P*<0.01 for comparison between stressed and unstressed males, and *P*<0.05 for comparison between stressed males and stressed females]. Spatial confinement also increased the frequency of spontaneous jerks across all groups (Suppl. Fig. 4B) [Main effect of stress: F(1,64)=12.71, *P*=0.0007], with male HZ mice exhibiting the highest frequency of body jerks regardless of stress exposure [Sex × genotype interaction: F(1,64)=20.39, *P*<0.0001]. Startle amplitude was higher in male mice [Main effect of sex: F(1,64)=9.20, *P*=0.003], but was not influenced by genotype, stress, or their interaction (Suppl. Fig. 4C). Finally, while spatial confinement reduced PPI across all groups [Main effect of stress: F(1,64)=4.06, *P*=0.048], a significant PPI deficit was observed only in unstressed *Celsr3* HZ males (Suppl. Fig. 4D) [Three-way genotype × stress × sex interaction, F(1,64)=4.03, *P*=0.049; *P*=0.005 for post-hoc comparisons between male HZ and WT].

To evaluate the predictive validity of *Celsr3* HZ mice as models of tic disorders, we assessed their behavioral responses to established Tourette Syndrome (TS) therapies. The D2 receptor antagonist haloperidol (0.25 mg/kg, administered IP 45 min before testing) normalized grooming behavior (Fig. 2A) [Genotype × treatment interaction: F(1,27)=11.61, *P*=0.002], head jerks (Fig. 2B) [Genotype × treatment interaction: F(1,28)=5.17, *P*=0.03], and PPI [Genotype × treatment interaction: F(1,28)=4.31, *P*=0.047], without affecting startle amplitude (Fig. 2C-D). The α2 receptor agonist clonidine (0.25 mg/kg, administered IP 30 min before testing) reduced stereotyped grooming (Fig. 2E) [Genotype × treatment interaction: F(1,28)=4.38, *P*=0.045] and tic-like behaviors (Fig. 2F) [Genotype × treatment interaction: F(1,28)=4.26, *P*=0.048]. Clonidine did not alter startle amplitude (Fig. 2G) but mitigated the PPI disruption in *Celsr3* HZ males (Fig. 2H) [Genotype × treatment interaction: F(1,28)=4.94, *P*=0.03]. The recently approved D1 receptor antagonist ecopipam^30^ (0.5–1.0 mg/kg, administered IP 20 min before testing) decreased spontaneous grooming (Fig. 2I) [Genotype × treatment interaction: F(2,42)=3.41, *P*=0.042] and head jerks (Fig. 2J) [Genotype × treatment interaction: F(2,42)=12.25, *P*<0.0001]. Ecopipam also improved the PPI deficits observed in *Celsr3* HZ males [Genotype × treatment interaction: F(2,42)=3.73, *P*=0.032], without affecting startle amplitude (Fig. 2K-L).

Finally, we tested the impact of the 5α-reductase inhibitor finasteride (10 mg/kg, administered IP 15 min before testing), as previous research has shown that this treatment reduces tic severity ^31^. As expected, finasteride reduced spontaneous grooming (Fig. 2M) [Genotype × treatment interaction: F(1,32)=6.50, *P*=0.02], decreased jerks (Fig. 2N) [Genotype × treatment interaction: F(1,32)=13.17, *P*=0.001], and reversed PPI deficits in *Celsr3* HZ mice [Genotype × treatment interaction: F(1,32)=12.75, *P*=0.001], without influencing startle amplitude (Fig. 2O-P).

**Transcriptomic analyses revealed significant alterations in RNA processing and cytoskeletal regulation in striatal tissues of *Celsr3* HZ mice.** Building on these findings, we concentrated our analyses on the molecular profile of the dorsal striatum. Immunostaining demonstrated that CELSR3 is expressed in this region, albeit at lower levels compared to the cortex, with HZ mice showing a significant reduction in expression across both the cortex and striatum (Suppl. Fig. 5A). Specifically, the analysis of CELSR3 levels in the striatum of HZ mice showed a significant decrease of approximately 43.75% compared to their WT littermates [F(1,22)=5.15, *P*=0.03] (Suppl. Fig. 5B).

Based on these results, we proceeded to analyze the transcriptomic profile of the striata of WT and HZ mice. Striata from 8 male animals per genotype were collected and processed for bulk transcriptomic analyses. All samples were sequenced for a total of 521.4 million (M) reads (average: 32.6 M; range: 27.2 M – 39.4 M). The average percentage mapping rate (uniquely mapped reads) was 82.7% (range: 40.3% - 89.3%). All the samples but one had a mapping rate larger than 79.9%. The sample with the lowest mapping rate (*WT_7*; mapping rate: 40.3%), appeared to be a significant outlier in the first round of PCA and was excluded from further analyses. After removing that sample, a second outlier was identified (sample: *HZ_3*) and removed from the dataset. No further outliers were identified in the third round of PCA, bringing the sample size for the final analysis to 7/genotype. Pearson's *r* revealed no significant correlation between RIN and the principal components 1 (*r* = 0.47; *P*= 0.090) and 2 (*r* = 0.04; P= 0.89). The comparison of 14 samples between *WT* and *HZ* showed 394 differentially expressed genes (DEGs) (Suppl. Table 1). Of these, 226 were upregulated, and 168 were downregulated (Suppl. Fig. 6A). The expression levels distribution in *HZ* and *WT* mice of the top 9 DEGs are represented in Suppl. Fig. 6B. As expected, *Celsr3* was among the top genes, showing a statistically significant downregulation in *HZ* (log_2_ FC = -0.817; FDR = 1.2E-03). Other top differentially expressed genes were: *Tmed2*, *Chaserr*, *Slc7a5*, *Rpph1*, and *Ap4s1* (Suppl. Table 1). The heatmap in Suppl. Fig. 6C, generated from the DEGs using the Euclidean distance with the Manhattan clustering method, showed clear-cut segregation of gene expression differences between *HZ* and *WT*. While no significant enrichment of cell-specific genes was observed (Suppl. Table 2), a significant enrichment of 18 distinct GO classes was found (Suppl. Fig. 6D). The top downregulated functional classes were “Structural Constituent of Cytoskeleton” (GO:0005200), “Structural Molecule Activity” (GO:0005198), and “Intercellular Bridge” (GO:0045171), whereas the top upregulated functional classes were “mRNA Processing” (GO:0006397), “RNA Splicing” (GO:0008380), and “RNA Splicing, via Transesterification Reactions” (GO:0000375).

**Spatial transcriptomic analyses revealed significant alterations in the extracellular matrix across the striatum**. To further investigate transcriptional variations across different regions of the striatum, detailed spatial transcriptomic analyses were conducted. Tissue slices were obtained from adult male WT and HZ mice, covering the area from -0.5 to +0.5 mm (AP). The spatial analysis identified 31 unique clusters (Suppl. Fig. 7A), seven of which were found within the striatum (specifically numbers 0, 3, 10, 21, 25, 27, and 30), all displaying enrichment for *Drd1* and *Drd2* (Suppl. Fig. 7B).

These clusters were categorized based on anatomical features and spatiomolecular markers^32^ (Fig. 3A-C, Suppl. Table 3): 1) the dorsolateral striatum, distinguished by high expression of *Nefm*, *Cnr1*, and *Gpr155*; 2) the dorsomedial striatum, notably enriched for *Crym*, *Col6a1*, and *Cpne6*; 3) the central striatum, strongly enriched in markers associated with white matter, such as *Mobp*, *Mag*, *Fth1*, *Mbp*, and *Mog*; 4) the ventral striatum, enriched in *Wfs1* and *Syt10*; 5) a cluster consisting of intermittent punctate areas with high enrichment for *Sst*, *Npy*, and *Nos1*, distinctive markers of somatostatin-NPY-NOS1-positive interneurons (SNNINs); 6) a cluster of intermittent puncta enriched with markers for cholinergic interneurons (CINs), such as *Chat*, *Slc17a8*, and *Ntrk1*; and 7) a cluster of sporadic puncta enriched in perivascular markers, including those for endothelial cells (*Vwf*, *Cldn5*) and pericytes (*Igf2*) (Suppl. Table 3).

The analysis across different regions revealed distinct pathway differences, highlighting unique biological functions, cellular components, molecular functions, and Reactome pathways in each area (Suppl. Tables 4-6).

Analyzing the global signal, we identified 233 DEGs, with 94 genes downregulated and 139 upregulated. Globally, key biological functions included “Negative Regulation of Blood Vessel Morphogenesis” (GO:2000181), “Cilium Movement” (GO:0003341), and “Negative Regulation of Angiogenesis” (GO:0016525). Structural differences were observed in cellular components such as the “Collagen-Containing Extracellular Matrix” (GO:0062023) and the “9+2 Motile Cilium” (GO:0097729) (Suppl. Table 4).Specifically, in terms of biological processes, there was a notable downregulation in processes such as “Negative Regulation of Blood Vessel Morphogenesis” (GO:2000181), “Negative Regulation of Angiogenesis” (GO:0016525), “Regulation of Mononuclear Cell Migration” (GO:0071675), “Adenylate Cyclase-Activating G Protein-Coupled Receptor Signaling Pathway” (GO:0007189), and “Cilium Movement” (GO:0003341). For cellular components, upregulation was observed in the “9+2 Motile Cilium” (GO:0097729) and “Cation Channel Complex” (GO:0034703), whereas there was a downregulation in components such as the “Collagen-Containing Extracellular Matrix” (GO:0062023) and “Endoplasmic Reticulum Lumen” (GO:0005788). Reactome pathway analysis revealed a downregulation in “Class B/2 (Secretin Family Receptors)” (R-HSA-373080), “Platelet Activation, Signaling, and Aggregation” (R-HSA-76002), and “GPCR Ligand Binding” (R-HSA-500792) (Fig. 3E and Suppl. Tables 5-6).

In the dorsolateral striatum, 169 DEGs were identified, with 104 genes upregulated, including *Krt8*, *Hmx3*, and *Cdh23*, while 65 genes such as *F5*, *Tekt2*, and *Adgb* were downregulated. In this region, biological functions such as “Negative Regulation of Angiogenesis” (GO:0016525) and “Macrophage Migration” (GO:1905517) were notably distinct. Differences in cellular components included the “Collagen-Containing Extracellular Matrix” (GO:0062023) and “Cell-Cell Junction” (GO:0005911), with molecular functions showing variations in “Calcium Ion Binding” (GO:0005509). Reactome pathways, such as “Collagen Chain Trimerization” (R-HSA-8948216) and “Collagen Formation” (R-HSA-1474290), also exhibited unique patterns (Suppl. Table 4) . Specifically, the dorsolateral striatum displayed increased activity in biological processes like "Neutrophil Chemotaxis" (GO:0030593), "Inflammatory Response" (GO:0006954), "Granulocyte Chemotaxis" (GO:0071621), “Neutrophil Migration” (GO:1990266), and "Negative Regulation of Blood Vessel Morphogenesis" (GO:2000181). Cellular components such as "Collagen-Containing Extracellular Matrix" (GO:0062023) and "Catenin Complex" (GO:0016342) were upregulated, whereas the expression of genes related to "Endoplasmic Reticulum Lumen" (GO:0005788), “Collagen-containing Extracellular Matrix” (GO: 0062023), and "Intracellular Organelle Lumen" (GO:0070013) were decreased. Molecular function analyses highlighted a significant upregulation in "G Protein-Coupled Receptor Activity" (GO:0004930), “Calcium Ion Binding” (GO:0005509), and “Chemoattractant Activity” (GO:0042056). Reactome analyses pointed out the downregulation in "Collagen Chain Trimerization" (R-HSA-8948216) and other collagen-related pathways (Fig. 3D-E and Suppl. Tables 5-6).

The dorsomedial striatum revealed 43 DEGs, with 30 upregulated genes, including *Crispdl2*, *St8sia6*, and *C1ql4*, and 13 downregulated genes such as *Adgb*, *Akap14*, and *Lox.* In this area, chemokine-related signaling was a notable biological function, with pathways such as the “Chemokine-Mediated Signaling Pathway” (GO:0070098) and processes related to extracellular matrix organization, including Reactome pathways like “Collagen Degradation” (R-HSA-1442490) and “Collagen Formation” (R-HSA-1474290). In the central striatum (CS), unique biological functions included “Cilium Movement” (GO:0003341) and “Inorganic Cation Transmembrane Transport” (GO:0098662), with cellular components like the “9+2 Motile Cilium” (GO:0097729) and the “Sperm Flagellum” (GO:0036126) showing distinct features (Suppl. Table 4). Specifically, in the dorsomedial striatum, biological processes related to "Chemokine-mediated Signaling Pathway” (GO:0070098) and “Cellular Response to Chemokine” (GO:1990869) were upregulated. Cellular components like "(9+2) Motile Cilium” (GO:0097729) showed downregulation. The molecular functions indicated heightened "CCR Chemokine Receptor Binding” (GO:0048020), “Chemokine Activity” (GO:0008009) and “Chemokine Receptor Binding” (GO:0042379). Reactome analysis highlighted upregulation in "Collagen Degradation” (R-HSA-1442490), "Degradation of Extracellular Matrix” (R-HSA-1474228), and “Activation of Matrix Metalloproteinases” (R-HSA-1592389) (Fig. 3D-E and Suppl. Tables 5-6).

In the central striatum, 143 DEGs were identified, with 97 genes like *Frmpd2*, *Cfap52*, and *Ttc29* upregulated, and 46 genes downregulated, including *Defb30*, *Prr29*, and *C1ql4*. This region was characterized by the increased activity in biological processes such as "Cilium Movement” (GO: 0003341), and cellular components such as “(9+2) Motile Cilium” (GO:0097729) (Fig. 3D and Suppl. Tables 4-6).

In the ventral striatum, 128 DEGs were detected, with 66 genes upregulated, such as *Mki67*, *Ppp1r1b*, and *Cxcl10*, while 62 genes, including *Zfp185*, *Msx1*, and *Igfbpl1*, were downregulated. This region displayed unique cellular components, notably the “Collagen-Containing Extracellular Matrix” (GO:0062023), and molecular functions linked to “Phosphatase Inhibitor Activity” (GO:0019212). In p-SNNINs, biological functions related to “Cilium Movement” (GO:0003341), “Cilium Movement Involved in Cell Motility” (GO:0060294), and “Blood Circulation” (GO:0008015) were prominent, emphasizing specific roles in cell motility and circulation (Suppl. Table 4). Specifically, the the ventral striatum exhibited upregulated expression of genes related to "Brain development” (GO: 0007420), and reductions in “Negative Regulation of Phospholipase Activity” (GO: 0010519) and “Collagen-Containing Extracellular Matrix" (GO:0062023) (Fig. 3D-E and Suppl. Tables 5-6).

Striatal SNNIN-enriched puncta exhibited 48 DEGs (33 upregulated and 15 downregulated). GO analyses found upregulations in “Cilium movement” (GO:0003341), “Sensory Perception” (GO: 0007600), and “T Cell Chemotaxis” (GO: 0010818) (Fig. 3D and Suppl. Tables 4-6).

Striatal puncta enriched in CINs showed only 7 DEGs (3 upregulated and 4 downregulated) between genotypes. Finally, no genotype differences were found with respect to the striatal puncta enriched in perivascular cells (Fig. 3E and Suppl. Tables 4-6).

**Single-nucleus transcriptomic analyses revealed cell-specific differences in gene expression in *Celsr3* HZ mice.** Given that the differential signature between the striata of *Celsr3* HZ and WT mice only pointed to broad functional alterations, we used single-nucleus (sn) transcriptomic methodologies to discern potential cell-specific differences in gene expression. SnRNA-seq was executed on the striatum derived from six adult male mice (n=3/genotype). Following quality control and normalization, we analyzed 48834 high-quality cells, of which 23706 were derived from WT and 25128 were from *Celsr3* HZ mice (Suppl. Table 7). Annotations utilizing recognized marker genes through SingleR pointed to several well-defined clusters corresponding to distinct cell types in the striatum, including 30193 neurons, 8775 oligodendrocytes, 5226 astrocytes, 2417 microglial cells, and 611 endothelial cells (Suppl. Table 7). We used marker genes to identify separate clusters corresponding to seven main cell populations, namely projection neurons, interneurons, astrocytes, oligodendrocytes, microglia, ependymal cells, and endothelial cells (Suppl. Figs. 8 and 9). We found 114 DEGs (14 upregulated and 100 downregulated) in the collective neuronal population from *Celsr3* HZ and WT mice. Comparisons between genotypes revealed that the clusters corresponding to projection neurons, interneurons, and microglia exhibited the most differences in gene expression (Suppl. Table 8 and Suppl. Fig. 8C). Probing deeper into specific neuronal clusters (Suppl. Fig. 10) characterized by established gene marker sets, a group corresponding to striatal projection neurons (SPNs) was identified, with 75 overall DEGs (10 upregulated and 65 downregulated). Notable among the 10 upregulated DEGs was *Ppp1r2*, a key regulator of DARPP-32 signaling of dopamine receptors. Gene ontology analyses of genes identified significant changes in several pathways, including the “Negative Regulation of Protein Phosphorylation” (GO:0001933) and “Positive Regulation of Glycolytic Process” (GO:0045821). In another cluster matching the characteristics of interneurons, 130 DEGs were found (43 upregulated and 87 downregulated), with gene-ontology alterations in “Axon Extension” (GO:0048846 and GO:0048675), “Axon Guidance” (GO:0007411), “Negative Chemotaxis” (GO:0050919) as well as “Glutamate Receptor Signaling Pathway” (GO:0007215), “Cell-Cell Adhesion via Plasma-Membrane Adhesion Molecules” (GO:0098742; GO:0007156 and GO:0007157). Comparative analyses of gene expression changes within microglial cells identified 92 DEGs (26 upregulated and 66 downregulated), with significant downregulations in several biological pathways, including “Synapse Pruning” (GO:0098883), “Cell Junction Disassembly” (GO:0150146), “Humoral Immune Response Mediated by Circulating Immunoglobulin” (GO:0002455), “Positive Regulation of Chemotaxis” (GO:0050921), and “Microglial Cell Activation” (GO:0001774). Taken together, these alterations point to subtle yet significant modifications of the interactions between SPNs, interneurons, and microglia within the striatum of *Celsr3* HZ mice. Conversely, oligodendrocytes, astrocytes, ependymal and endothelial cells showcased only modest transcriptomic deviations between genotypes, with mild upregulations in the positive regulation of transport and phagocytosis in both cell types (Suppl. Fig. 8C). These data suggest a lesser vulnerability of these cell types to the transcriptional impact of a partial *Celsr3* deficiency.

Given that these results pointed to subtle functional deficits in generic clusters of SPNs and interneurons, to gain more insight into the specific subpopulations of affected cells, we re-analyzed our snRNA-seq data using a Gamma-Poisson generalized linear model with a 0.4 clustering resolution for Seurat (Suppl. Figs. 11-13 and Suppl. Tables 9-12), since this platform is particularly useful in capturing alterations in transcriptomic profiles of specific cell populations within single cell databases. Identification of specific clusters for SPNs led to the annotation of six separate clusters (as identified by high *Rgs9* expression) (Fig. 4A-B). Using curated lists of marker genes for different striatal populations, we found four clusters matching canonic SPN identifiers, such as *Foxo1*. These four clusters were classified in relation to their positivity to *Drd1* (indicative of direct pathway) or *Drd2* (indicative of indirect pathway) as well as their location in the matrix or striosomes (patches). Thus, we identified D_1_-positive matrisomal SPNs (mD1-SPNs; enriched in *Drd1, Tac1,* and high expression of *Epha4;* 1012 DEGs), D_1_-positive striosomal SPNs (sD1-SPNs; enriched in *Drd1, Tac1, Sema5b, Oprm1,* and *Pdyn*; 296 DEGs), D_2_-positive matrisomal SPNs (mD2-SPNs; enriched in *Drd2*, *Adora2a, Penk,* and high expression of *Epha4;* 807 DEGs), and D_2_-positive striosomal SPNs (sD2-SPNs; enriched in *Drd2, Adora2a, Penk, Sema5b,* and high expression of *Foxp2*; 90 DEGs) (Suppl. Tables 9-12).

In addition, we identified a clusters with identifiers overlapping with those described for “exopatch” neurons (e-SPNs), such as *Col11a1, Otof, Casz1,* and *Tshz1.* Notably, this cluster featured high transcript levels of the genes encoding *Drd1* and *Drd3*, encoding the dopamine receptors D_1_ and D_3_ (Suppl. Tables 9-12).

Using the same procedure, we identified five clusters of interneurons featuring unique identifiers for cholinergic interneurons (*Chat*), GABAergic parvalbumin-positive interneurons (*Pvalb*, 57 DEGs), GABAergic calretinin-positive interneurons (*Calb2*, 48 DEGs), GABAergic somatostatin-NPY-nitric oxidase synthase 1 positive interneurons *(Sst, Npy, Nos1*, 29 DEGs), and GABAergic NPY-neurogliaform interneurons (enriched for *Npy*, but not *Sst* and *Nos1*, 1 DEG) (Fig. 4A-B).

In the global comparison between WT and *Celsr3* HZ mice, upregulated pathways revealed enhanced activity in neuronal development and signaling, including "Neuron Projection Development" (GO:0031175), "Phosphatidylinositol Binding" (GO:0035091), and "Signaling By Rho GTPases" (R-HSA-194315) (Fig 5C and Suppl. Table 11). Key structural components such as "Neuron Projection" (GO:0043005) and "Axon" (GO:0030424) were also enriched, indicating changes in cellular architecture (Fig. 5D and Suppl. Table 12). Conversely, downregulated pathways highlighted a reduction in metabolic and translational processes, such as "Energy Derivation By Oxidation Of Organic Compounds" (GO:0015980) and "Cytoplasmic Translation" (GO:0002181), along with decreased activity in "Mitochondrial Respiratory Chain Complex IV" (GO:0005751) and "Cytosolic Small Ribosomal Subunit" (GO:0022627), suggesting an overall decline in energy metabolism and protein synthesis in *Celsr3* HZ mice (Suppl. Fig. 13A and Suppl. Table 12).

In the mD1-SPNs cluster, upregulated pathways revealed increased activity in neuronal growth and regulatory processes, including "Neuron Development" (GO:0048666), "Regulation Of Neuron Projection Development" (GO:0010975), and "Positive Regulation Of Autophagy" (GO:0010508) (Fig 5C and Suppl. Table 11). Cellular components with enhanced expression included "Neuron Projection" (GO:0043005), "Asymmetric Synapse" (GO:0032279), and "Postsynaptic Density" (GO:0014069), indicating structural changes in synaptic architecture. Molecular functions like "Phosphatidylinositol Binding" (GO:0035091), "Zinc Ion Binding" (GO:0008270), and "Transition Metal Ion Binding" (GO:0046914) were also upregulated, suggesting shifts in receptor activity and ion interactions (Fig 5D and Suppl. Table 11). Enriched Reactome pathways included "Signaling By Nuclear Receptors" (R-HSA-9006931), "Chromatin Modifying Enzymes" (R-HSA-3247509), and "ESR-mediated Signaling" (R-HSA-8939211), collectively indicating increased transcriptional and chromatin remodeling activity (Suppl. Table 11).

In the mD2-SPNs cluster, upregulated pathways highlighted increased activity in neuron structure and synaptic processes, including "Neuron Projection Development" (GO:0031175), "Neuron Development" (GO:0048666), and "Clathrin Coat Disassembly" (GO:0072318) (Fig. 5C and Suppl. Table 11). Enriched cellular components included "Neuron Projection" (GO:0043005), "Postsynaptic Density" (GO:0014069), and "Asymmetric Synapse" (GO:0032279), suggesting changes in synaptic architecture (Fig. 5D and Suppl. Table 11). Molecular functions with increased expression were "PDZ Domain Binding" (GO:0030165), "Phosphatidylinositol Binding" (GO:0035091), and "Glutamate Receptor Binding" (GO:0035254), indicating shifts in receptor interactions and signal modulation (Fig. 5E and Suppl. Table 11). Reactome pathways upregulated in this cluster included "Signaling By NTRK1 (TRKA)" (R-HSA-187037), "Intracellular Signaling By Second Messengers" (R-HSA-9006925), and "Signaling By NTRKs" (R-HSA-166520), pointing to active intracellular signaling (Suppl. Table 11). Conversely, downregulated Reactome pathways included "Peptide Chain Elongation" (R-HSA-156902), "Nonsense Mediated Decay (NMD) Enhanced By Exon Junction Complex (EJC)" (R-HSA-975957), and "Selenocysteine Synthesis" (R-HSA-2408557), suggesting a reduction in translational and protein synthesis-related processes in this cluster (Suppl. Table 12).

In the sD1-SPNs cluster, upregulated pathways emphasized enhanced signaling and synaptic processes, including "Retinoic Acid Receptor Signaling Pathway" (GO:0048384), "Chemical Synaptic Transmission" (GO:0007268), and "Positive Regulation Of Cell Projection Organization" (GO:0031346) (Fig. 5C and Suppl. Table 11). The cellular component "Neuron Projection" (GO:0043005) was notably enriched, indicating changes in neuronal projection structures (Fig. 5D and Suppl. Table 11). Molecular functions with increased expression included "Amino Acid Binding" (GO:0016597), "Adenylate Cyclase Regulator Activity" (GO:0010854), and "Leucine Binding" (GO:0070728), suggesting shifts in receptor and regulatory activities (Fig. 5E and Suppl. Table 11). Enriched Reactome pathways were "Signaling By Rho GTPases" (R-HSA-194315), "Signaling By Rho GTPases, Miro GTPases And RHOBTB3" (R-HSA-9716542), and "Muscle Contraction" (R-HSA-397014), collectively pointing to active roles in cellular signaling and structural regulation within this cluster (Suppl. Table 11).

In the sD2-SPNs cluster, the upregulated biological process included "Protein Targeting To Vacuole Involved In Autophagy" (GO:0071211) (Fig. 5C and Suppl. Table 11). Conversely, the downregulated cellular component was the "beta-catenin-TCF Complex" (GO:1990907) (Suppl. Fig. 13B and Suppl. Table 12).

In the e-SPNs cluster, upregulated pathways included biological processes "Sodium Ion Transport" (GO:0006814), "Detection Of Stimulus Involved In Sensory Perception Of Pain" (GO:0062149), and "Metal Ion Transport" (GO:0030001) (Fig. 5C and Suppl. Table 11). Enriched cellular components were the "Sarcoglycan Complex" (GO:0016012) and "Dystroglycan Complex" (GO:0016011) (Fig. 5D and Suppl. Table 11).

In the CRINs cluster, upregulated pathways included biological processes "Regulation Of Synapse Assembly" (GO:0051963), "Positive Regulation Of Cell Projection Organization" (GO:0031346), and "Enzyme-Linked Receptor Protein Signaling Pathway" (GO:0007167), highlighting enhanced activities in synapse formation and cell projection organization (Fig. 5C and Suppl. Table 11). Enriched Reactome pathways involved "Receptor-type Tyrosine-Protein Phosphatases" (R-HSA-388844), "Neuronal System" (R-HSA-112316), and "Protein-protein Interactions At Synapses" (R-HSA-6794362), suggesting active synaptic signaling and receptor interactions (Suppl. Table 11). Conversely, downregulated pathways in biological processes included "Regulation Of Monoatomic Ion Transmembrane Transporter Activity" (GO:0032412), "Regulation Of Synaptic Transmission, Glutamatergic" (GO:0051966), and "Positive Regulation Of Long-Term Synaptic Potentiation" (GO:1900273), pointing to reduced ion transport and glutamatergic transmission (Suppl. Fig. 13A and Suppl. Table 12). Downregulated cellular components were "Neuron Projection" (GO:0043005) and "Dendrite" (GO:0030425) (Suppl. Fig. 13B and Suppl. Table 12), while the molecular function "Cyclic Nucleotide Binding" (GO:0030551) was also decreased, indicating shifts in synaptic and ion transport mechanisms in this cluster (Suppl. Table 12).

In the PVINs cluster, upregulated pathways included biological processes "Sprouting Angiogenesis" (GO:0002040), "Anterograde Trans-Synaptic Signaling" (GO:0098916), and "Chemical Synaptic Transmission" (GO:0007268), indicating enhanced activity in vascular growth, synaptic signaling, and neurotransmission (Fig. 5C and Suppl. Table 11). Enriched cellular components were "Postsynaptic Density Membrane" (GO:0098839), "Postsynaptic Specialization Membrane" (GO:0099634), and "Synaptic Membrane" (GO:0097060), highlighting adaptations in synaptic structure (Fig. 5D and Suppl. Table 11). Molecular functions with increased activity included "Transmitter-Gated Monoatomic Ion Channel Activity" (GO:0022824), "Ligand-Gated Monoatomic Ion Channel Activity Involved In Regulation Of Presynaptic Membrane Potential" (GO:0099507), and "Neurotransmitter Receptor Activity Involved In Regulation Of Postsynaptic Membrane Potential" (GO:0099529), indicating upregulation in ion channel and receptor functions for membrane potential regulation (Fig. 5E and Suppl. Table 11). Reactome pathways enriched were "L1CAM Interactions" (R-HSA-373760), "Axon Guidance" (R-HSA-422475), and "Nervous System Development" (R-HSA-9675108), emphasizing enhanced roles in neuronal development and signaling within this cluster (Suppl. Table 11).

In the SNNINs cluster, upregulated pathways included biological processes "Protein Localization To Microtubule" (GO:0035372), "Protein Localization To Microtubule Cytoskeleton" (GO:0072698), and "Regulation Of Leukocyte Chemotaxis" (GO:0002688), indicating enhanced activity in protein positioning related to the microtubule cytoskeleton and immune cell movement (Fig. 5C and Suppl. Table 11). Conversely, the downregulated biological process was "Purine-Containing Compound Metabolic Process" (GO:0072521), suggesting a reduction in pathways associated with purine metabolism within this cluster (Suppl. Fig. 13A and Suppl. Table 12).

**Antagonism of D_3_ dopamine receptors reduces grooming and tic-like behaviors in *Celsr3* HZ mice.** Building on this evidence, we compiled a list of top DEGs for each neuronal cluster, focusing on those present in at least two cell types. (Fig. 5A). This list included 61 genes, such as serine-threonine kinases (*Sik2* and *Sik3*), key neurotrophin signaling molecules (including *Ntrk2*, encoding the NT3 growth factor receptor, and its signaling effector *Shc3*), myelin-related factors like *Mbp* (encoding myelin basic protein) and *Mobp* (myelin-associated oligodendrocyte basic protein), as well as thyroid hormone-binding targets such as *Ttr* (transthyretin) and *Thrb* (thyroid hormone receptor beta). Additionally, most cell types in *Celsr3* HZ mice showed downregulation of *Etv5* (encoding a transcription factor) and *Rpl29* (encoding a ribosomal protein). Notably, *Drd3* (encoding the D_3_ dopamine receptor) exhibited significant upregulation in sD1-SPNs and a simultaneous reduction in CRINs (Fig. 5A). These findings, along with previous reports indicating that *Celsr3* mutants exhibit increased dopamine release in the striatum ^33^, led us to examine D_3_ dopamine receptors in the nigrostriatal system using dopamine transporter (DAT), a key marker of dopaminergic fibers, alongside D_3_ receptor immunofluorescence and RNAscope^TM^ *in situ* hybridization (Fig. 6B-I). Leveraging RNAscope^TM^, we identified an increase in the mRNA levels of D_3_ receptors in the striatum [Fig. 6D, F(1,10)=5.44, P=0.04]. Furthermore, as depicted in Fig. 6E, we identified an increase in striatal cells positive for both D_1_ and D_3_ mRNAs [F(1,10)=6.44, P=0.03]. However, while *Celsr3* HZ mice showed an increase in D_3_ receptors also at protein level [Fig.6H, F(1,27)=6.46, P=0.02], there was a decrease in D_3_ presynaptic receptors in dopaminergic fibers, as indicated by the coexpression of D_3_ and DAT [Fig. 6I, F(1,27)=17.89, P<0.001]. In the substantia nigra *pars compacta*, *Celsr3* HZ mice showed a reduction in DAT immunofluorescence (F(1,22)=6.57, P=0.02] while overall D_3_ receptor expression and D_3_ levels in DAT-positive neurons remained consistent across genotypes (Suppl. Fig. 14).

Based on these results, we hypothesized that the alterations in striatal D_3_ receptor expression may contribute to the behavioral phenotype observed in HZ mice. Thus, we tested the effects on the spontaneous behavior of *Celsr3* male mice of two selective D_3_ drugs: the agonist PD-128907 (5 μg/kg, injected IP 10 min before testing) and the antagonist SB-277011A (10 mg/kg, injected IP 20 min before testing). PD-128907 dramatically increased grooming stereotypies in HZ, but not WT mice (Fig. 6A) [genotype × treatment interaction F(1,37)= 5.36, P=0.026]; furthermore, this compound increased tic-like responses in HZ mice (Fig. 6B) [genotype × treatment interaction F(1,34)=5.237, P=0.028]. In contrast, the D_3_ receptor agonist did not significantly impact startle amplitude and PPI (Fig. 6C-D). Notably, the effects of SB-288011A were opposed to those of PD-128907; indeed, this compound significantly reduced grooming in HZ mice without eliciting any effect in WT controls (Fig. 6E) [genotype × treatment interaction F(1,44)=8.29, P=0.006]; furthermore, this drug significantly reduced jerks in HZ mice (Fig. 6F) [genotype × treatment interaction F(1,44)=10.69, P=0.002], without impacting the startle amplitude or the PPI (Fig. 6G-H). Cell-specific changes in D_3_ expression seem to be implicated in TS-related behaviors, except for sensorimotor gating deficits.

**SUPPLEMENTAL FIGURES**

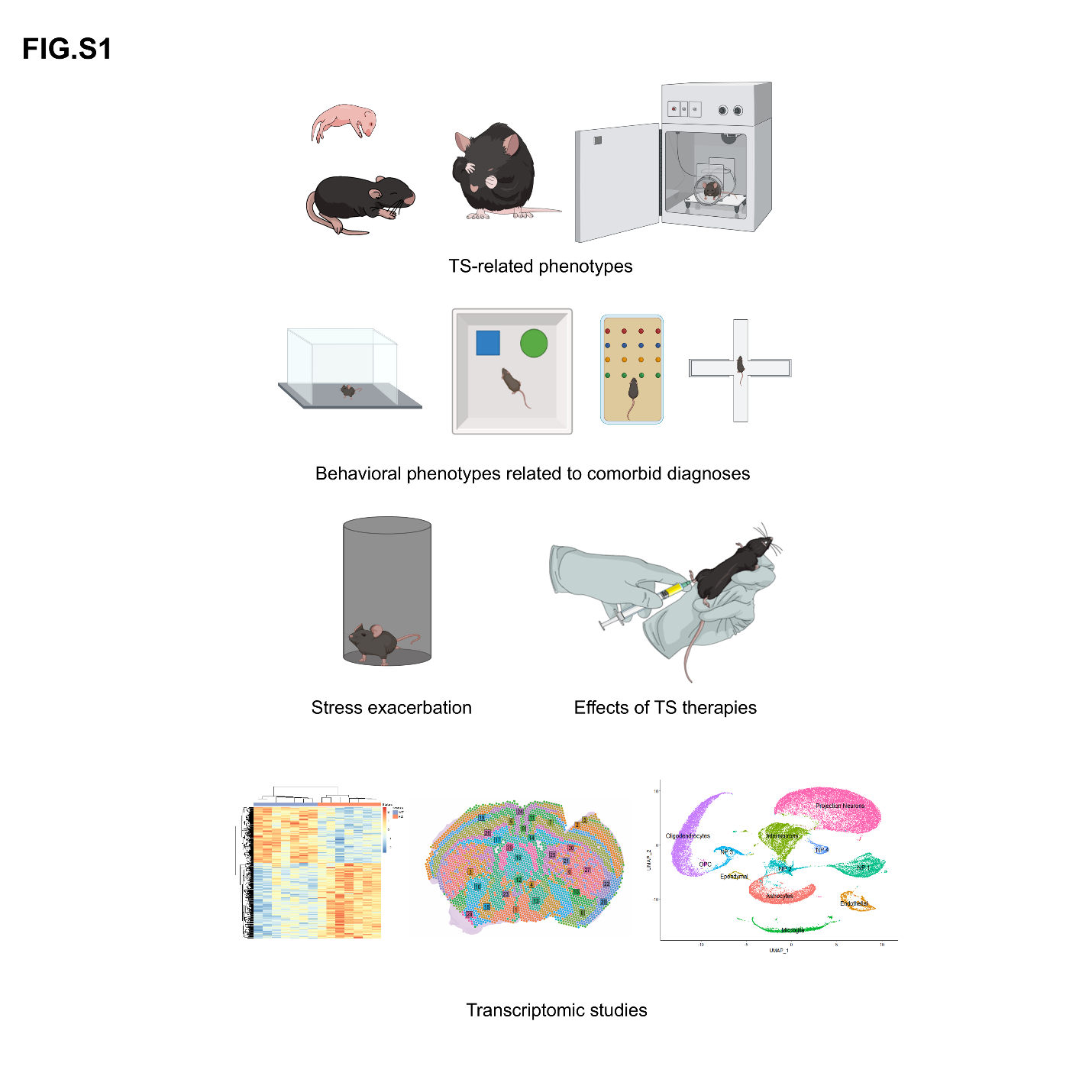

**Suppl. Fig. 1. Schematic representation of the procedures used in the present study.** The face validity of *Celsr3* heterozygous (HZ) mice with respect to tic disorders and related behavioral deficits was assessed at different developmental stages (neonates, adolescents, and adults, using different cohorts of mice to avoid carryover stress), in comparison with wild-type (WT) littermates. Separate groups of juvenile mice were tested to measure behavioral responses related to common comorbid diagnoses, such as anxiety and obsessive-compulsive disorder. The effects of acute stress and pharmacological treatments were also tested in juvenile mice to determine the predictive validity of *Celsr3* HZ mice. Following these analyses, we conducted bulk, spatial, and single-nucleus transcriptomic profiling of the striatum of *Celsr3* HZ and WT mice to identify neurobiological aberrances associated with Celsr3 deficiency.

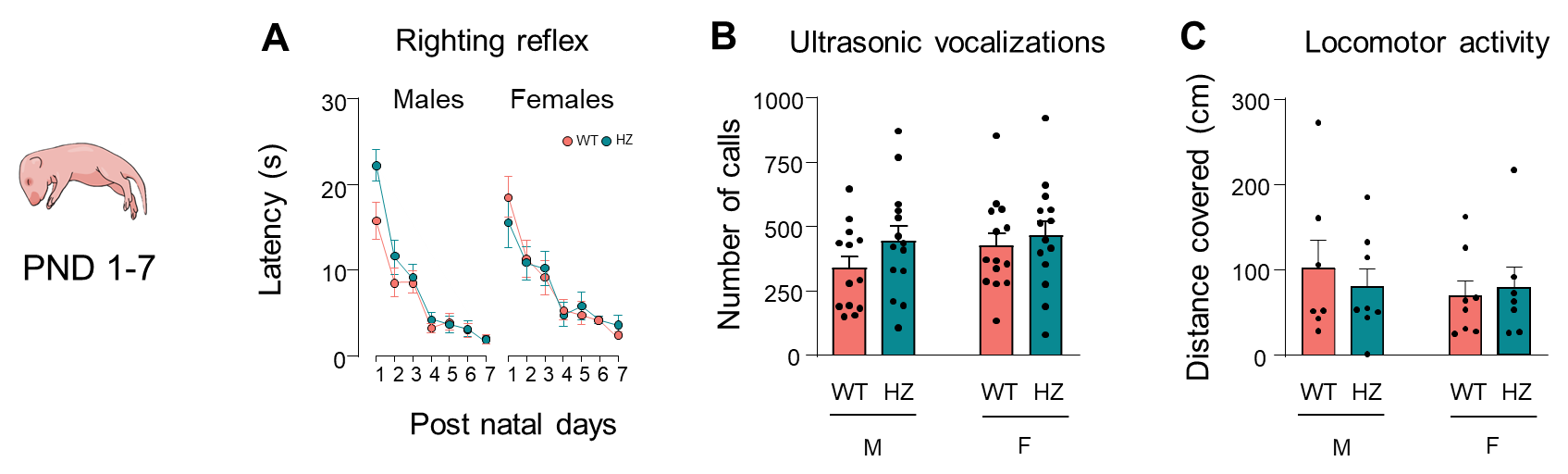

**Suppl. Fig. 2. *Celsr3* heterozygous (HZ) and wild-type (WT) littermates do not show any overt behavioral abnormalities between postnatal day (PND) 1 and 7.** (A) Righting reflex was equivalent between the two genotypes in both males and females throughout the first week of postnatal life. (B) Analyses of ultrasonic vocalizations (20-120 kHz) on PND 6 showed no differences between sexes or genotypes. (C) Locomotor activity was monitored on PND 7, and the total distance traveled was similar among groups. All data are shown as means ± SEM. WT groups are indicated in coral and HZ groups in teal (n= 15 female WT, 13 male WT, 16 female HZ, and 22 male HZ). No more than 1 pup/sex/genotype/litter was used to avoid litter effects. Data in panel A were analyzed by three-way, repeated-measure ANOVA, with time, sex, and genotype as factors. Data in panels B and C were analyzed by two-way ANOVA, with sex and genotype as factors. M, males; F, females.

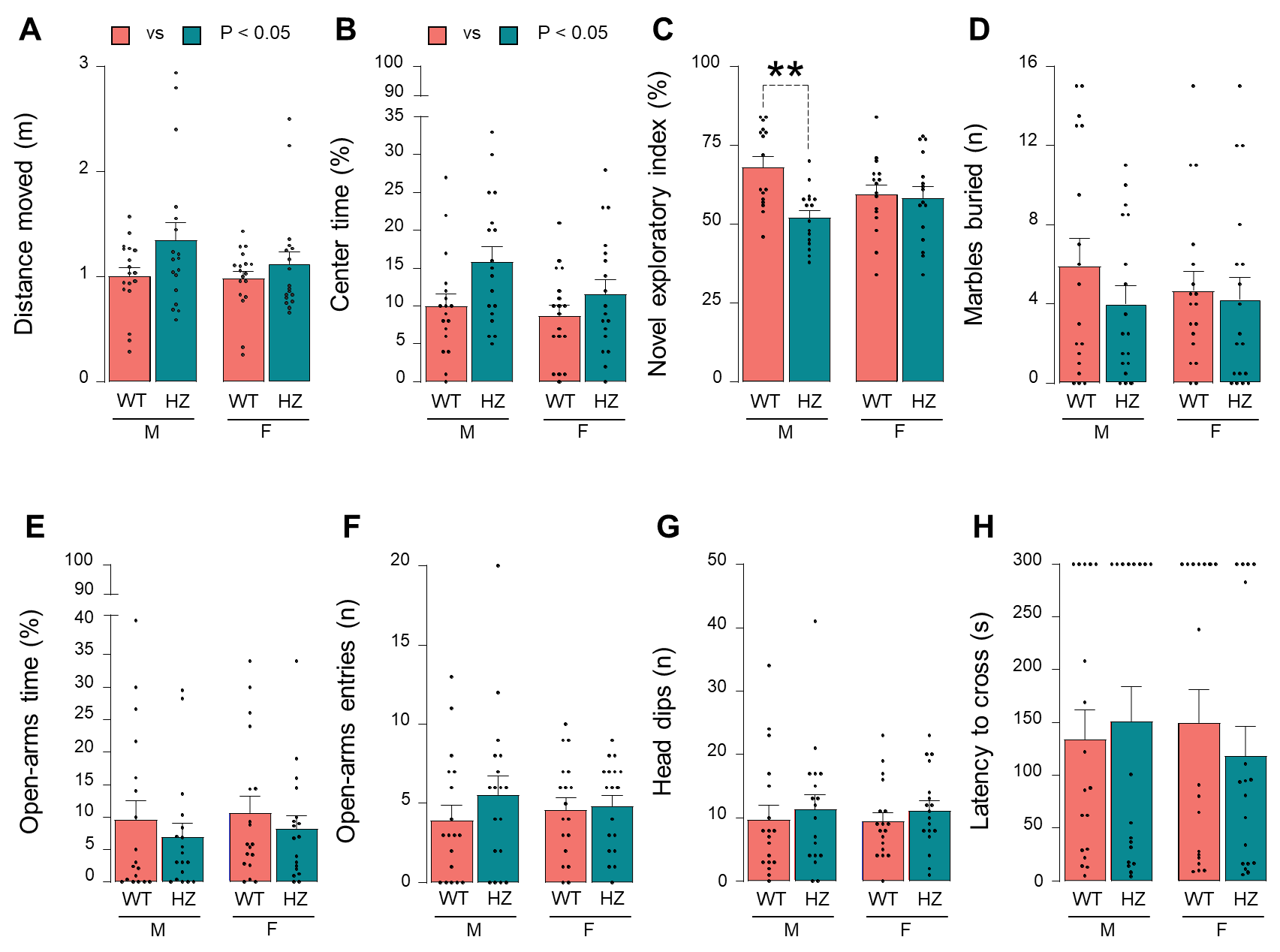
**Suppl. Fig. 3**. ***Celsr3* heterozygous (HZ) mice show comorbid hyperactivity and attentional deficits.** (A-B) Locomotor activity was monitored in a normal cage for 10 minutes. Evaluation of the average distance traveled (A) and the time spent in the center (B) showed a main effect for genotype, highlighting a hyperactive phenotype for *Celsr3* HZ mice. (C) Novel object recognition tests revealed a significant deficit in HZ males as compared to their wild-type (WT) counterparts (**, P<0.01). (D-H) No other differences between groups were identified independently of the paradigm used, whether it was to test compulsivity (marble burying, D), anxiety (elevated plus maze, E-G), or impulsivity (wire-beam elevated bridge test, H). Data are represented as means ± SEM with individual points. Data were analyzed with two-way ANOVA with genotype and sex as factors, followed by post-hoc analyses with Tukey´s correction (n=18/group, except for panel C, n=16/group). WT are indicated in coral and HZ in teal. M, males; F, females.

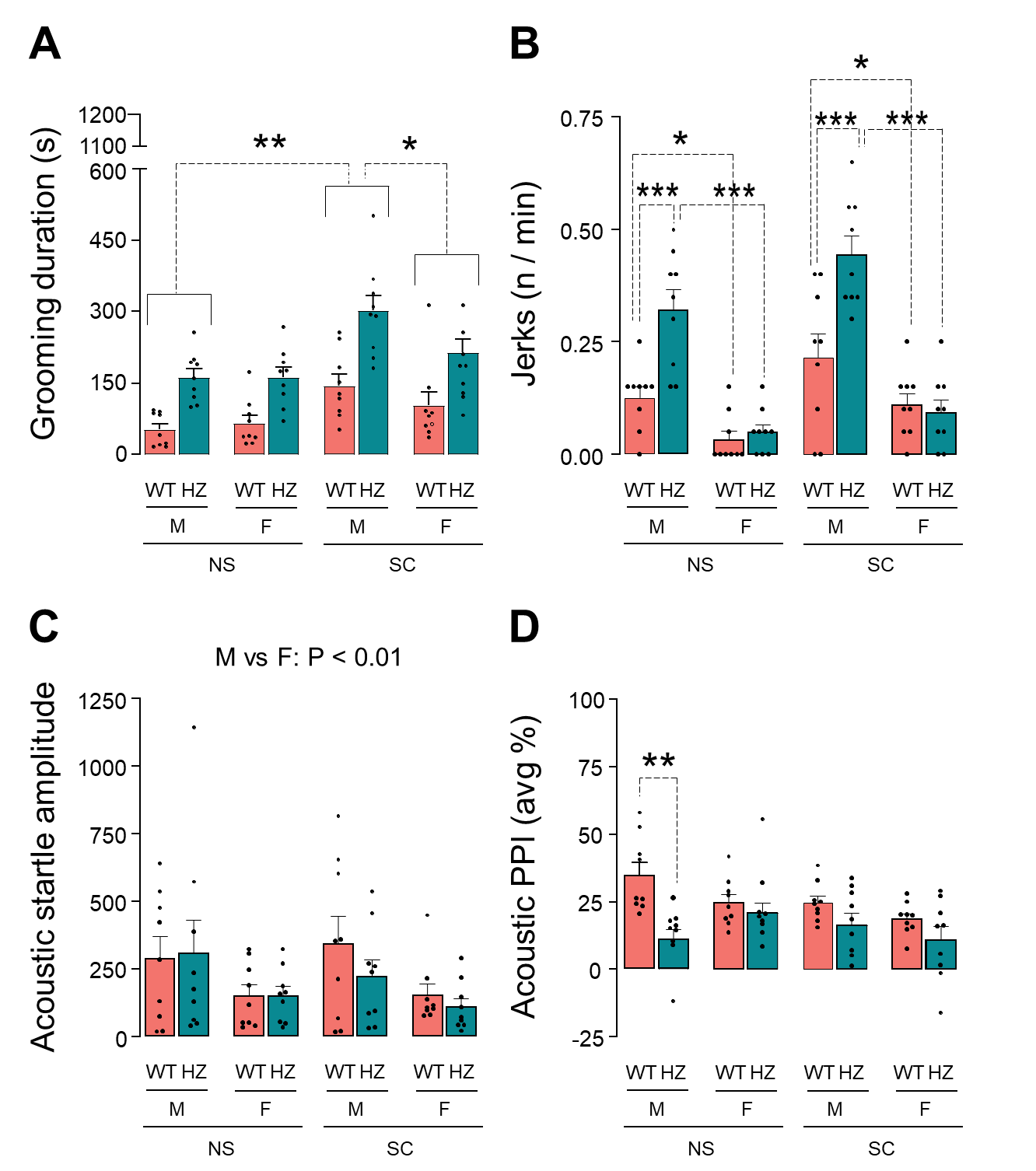

**Suppl. Fig. 4. Acute stress affects Tourette syndrome (TS)-related phenotypes differently in *Celsr3* heterozygous (HZ) male and female mice.** The effect of acute stress was studied by analyzing the behavior of naïve animals (NS, not stressed) with respect to animals exposed to spatial confinement (SC). SC elicited a significant increase in (A) grooming duration in both HZ and wild-type (WT) mice. Grooming duration was higher in males exposed to SC, compared to NS males and to SC females, irrespective of genotype. (B) The frequency of jerks depended on genotype and sex, but it was irrespective of stress. While *Celsr3* HZ males showed the highest frequency, female WT mice showed the lowest frequency of jerks. (C) The amplitude of acoustic startle reflex was higher in males than females, but no effects were detected depending on the genotype. (D) A significant deficit of prepulse inhibition (PPI) was detected only in HZ males not exposed to stress (NS), in comparison with NS WT males. PPI values were averaged across the three prepulse loudness levels. All data are shown as means ± SEM, with WT groups indicated in coral and HZ groups in teal (n= 9/group). M, males; F, females. All data were analyzed with three-way ANOVA with sex, genotype, and stress condition as factors. *, P<0.05, **, P<0.01, and ***, P<0.001 for post-hoc comparisons (indicated by brackets).

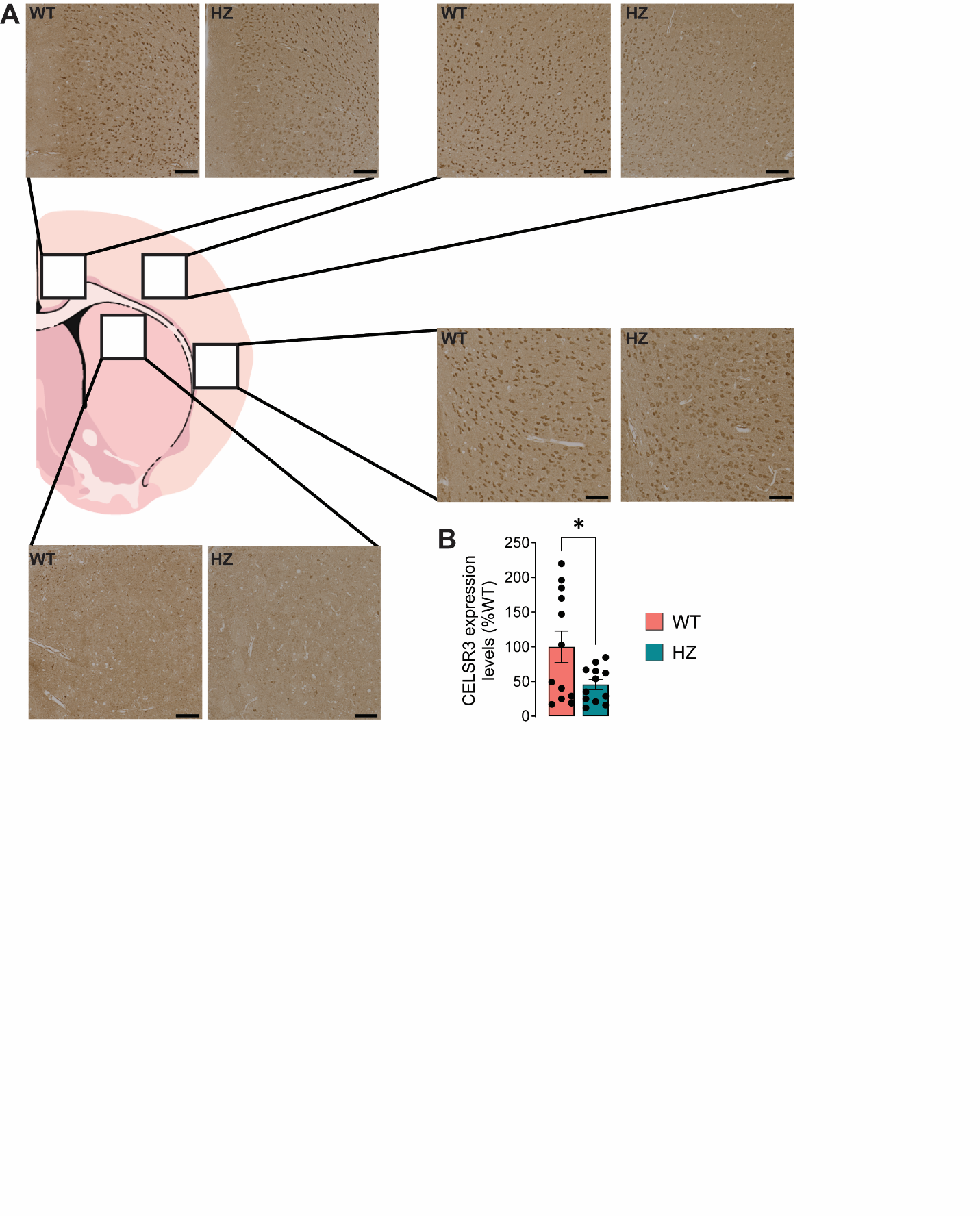
**Suppl. Fig. 5: CELSR3 distribution and expression in experimental animals.** (A) Representative images depicting immunohistochemistry analyses of CELSR3 expression in the cortex and striatum of wild-type (WT) and *Celsr3* heterozygous (HZ) animals. (B) Semi-quantitative analysis, performed using QuPath software (0.5.1), illustrates the reduction of CELSR3 staining in the striatum of *Celsr3* HZ animals compared to their WT counterparts. Data are presented as mean ± SEM (n=3/group). *, p<0.05.

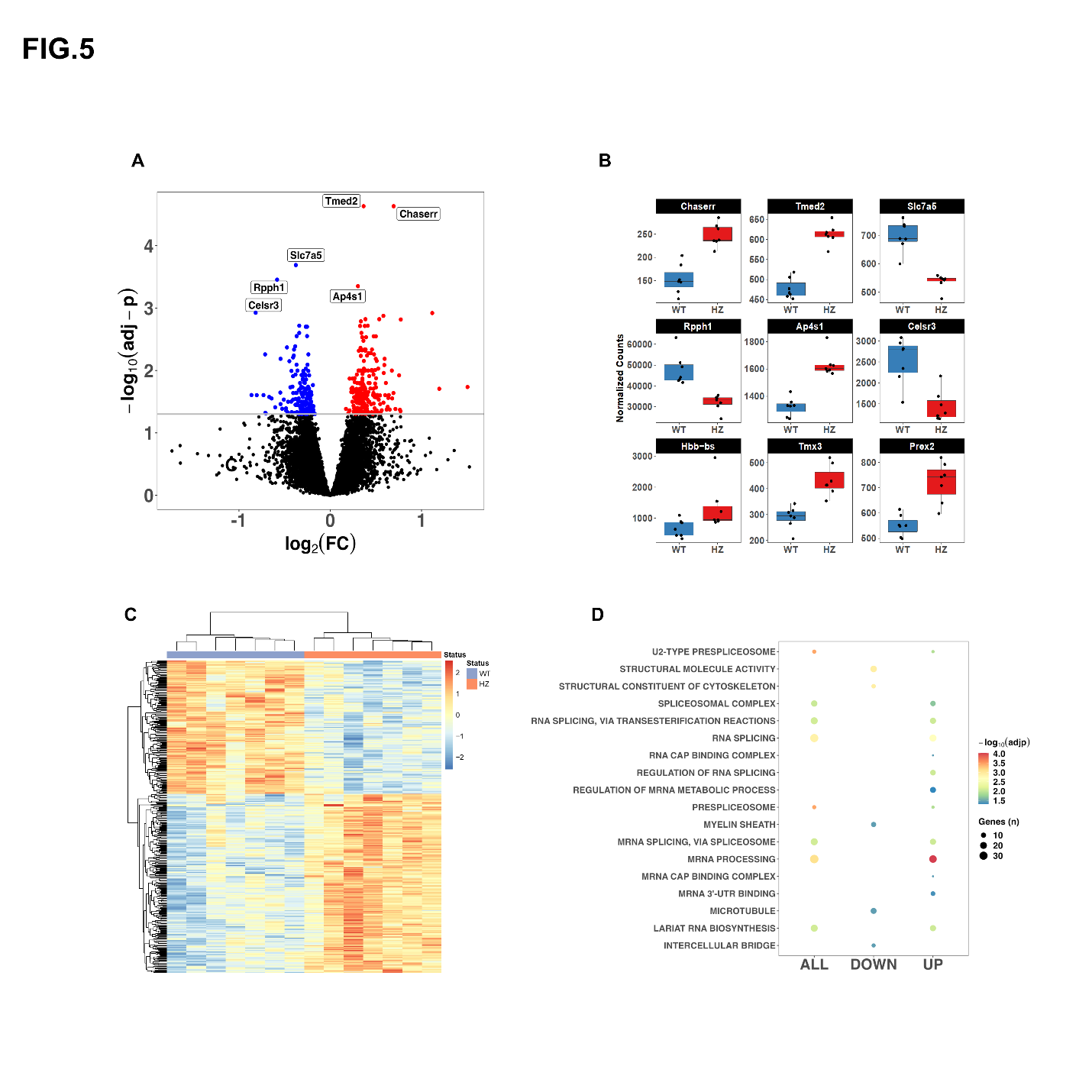

**Suppl. Fig. 6. Transcriptomic analyses of bulk striatal tissue of *Celsr3* heterozygous (HZ) and wild-type (WT) mice reveal differences in key metabolic processes.** (A) Volcano plot showing the differentially expressed genes (DEGs) between the *Celsr3* heterozygous (HZ, n=8) and wild-type (WT, n=8) striata. The top 6 DEGs are labeled. (B) Box plots showing the top 9 DEGs (HZ in red and WT in blue). (C) Heatmap generated with the Euclidean distance and Manhattan clustering method showing all the DEGs (FDR<0.05). (D) The results of gene ontology analyses using all the DEGs (all), only the downregulated (down), or only the upregulated (up) DEGs revealed that HZ mice showed differences in key processes, including the regulation of mRNA processing and metabolism, as well as microtubule regulation.

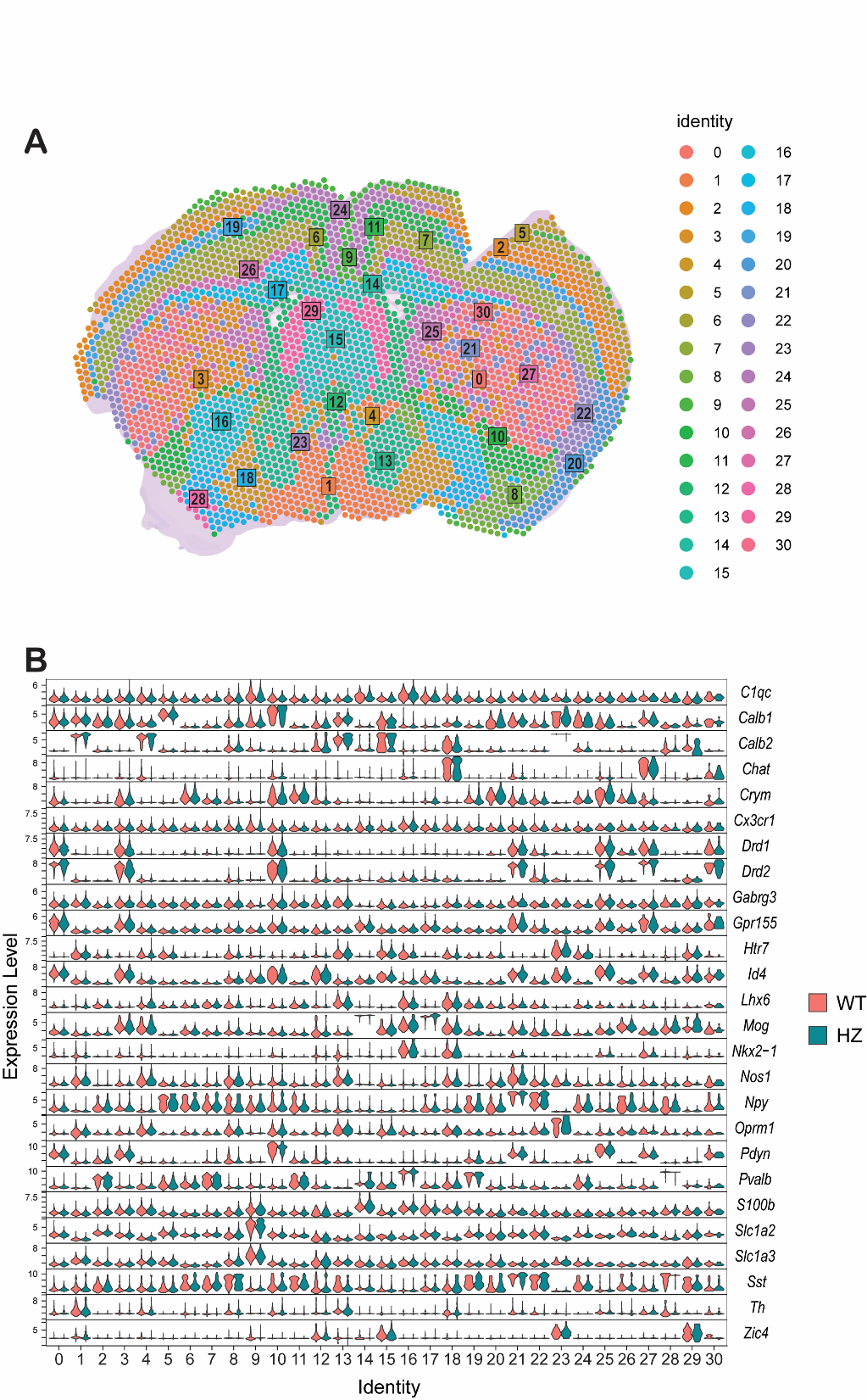

**Suppl. Fig. 7**. **Spatial transcriptomic analysis of gene expression in the brain of wild-type (WT) and *Celsr3* heterozygous (HZ) animals.** (A) Spatial distribution map representing distinct transcriptional identities across different brain regions. Each dot corresponds to a spatial barcode (identifying a 50 µm^2^ area), with regions color-coded based on identity clusters (0-30). (B) Violin plots showing the expression levels of selected genes across different spatial identities for WT and *Celsr3* HZ animals. The y-axis represents expression levels, and the x-axis indicates spatial identities corresponding to the clusters in panel (A).

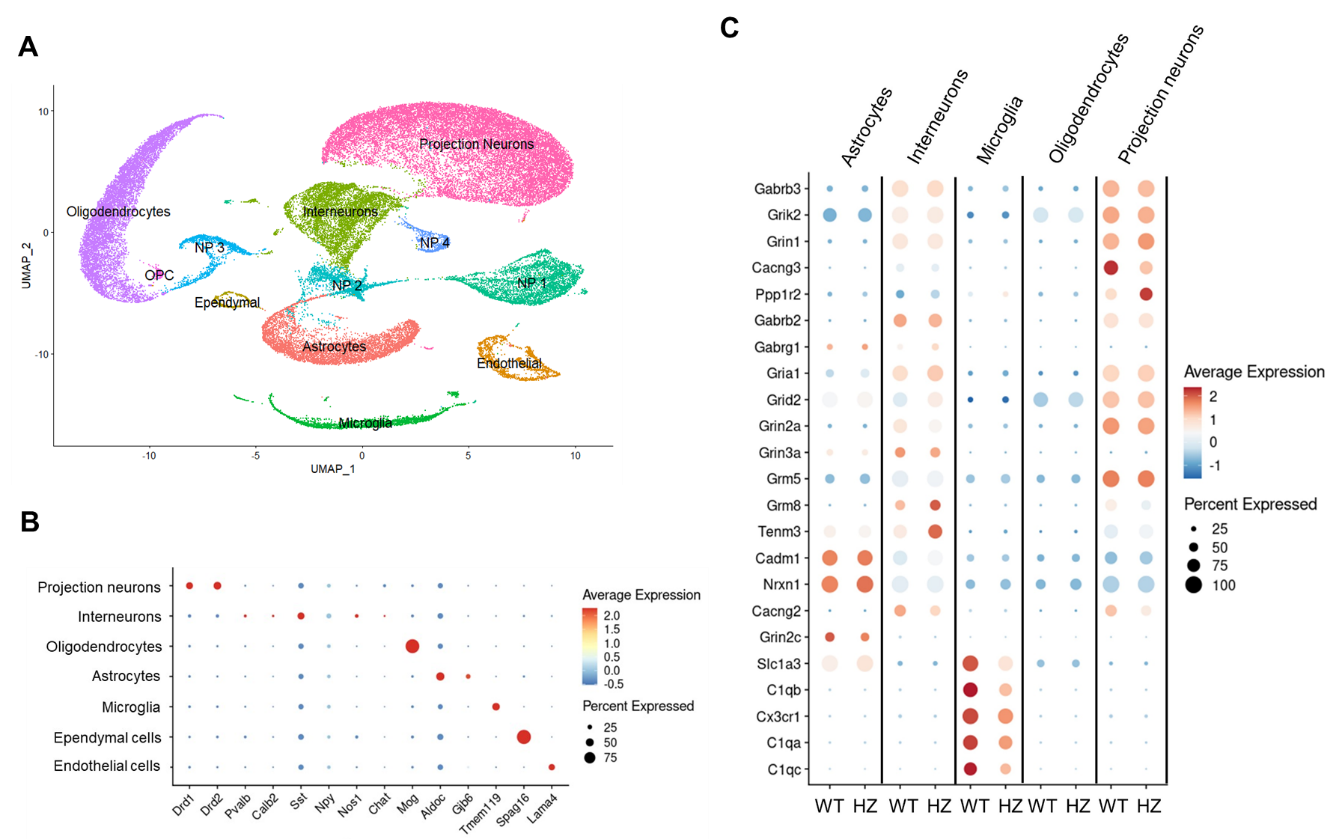
**Suppl. Fig. 8**. **Single-nucleus transcriptomic analyses of striatal tissues of *Celsr3* heterozygous (HZ) and wild-type (WT) mice reveal most differences in neuron and microglia clusters.** (A) Uniform Manifold Approximation and Projection (UMAP) of 48,834 nuclei annotated by major cell types, including astrocytes, oligodendrocytes, microglia, projection neurons, interneurons, progenitor cells (OPC), and other neuronal populations (NP 1-4). (B) Annotation of major clusters by select genes characteristic for major cell types. *Drd1* and *Drd2* expression is expected in projection neurons; *Pvalb*, *Calb2*, *Sst*, *Npy*, *Nos1*, and *Chat* as representative of several types of interneuron populations; *Mog* for mature oligodendrocytes; *Aldoc* and *Gjb6* for astrocytes; *Tmem119* for microglia; *Spag16* for ependymal cells; and *Lama4* for endothelial cells. (C) Dotplot of differential expression between wild-type (WT) and *Celsr3* heterozygotes (HZ). Decreased expression is observed in several microglial-expressed genes related, some modest alterations in astrocytes, almost no changes in oligodendrocytes, but many alterations in glutamatergic and GABAergic pathways in the projection neurons and interneuron populations.

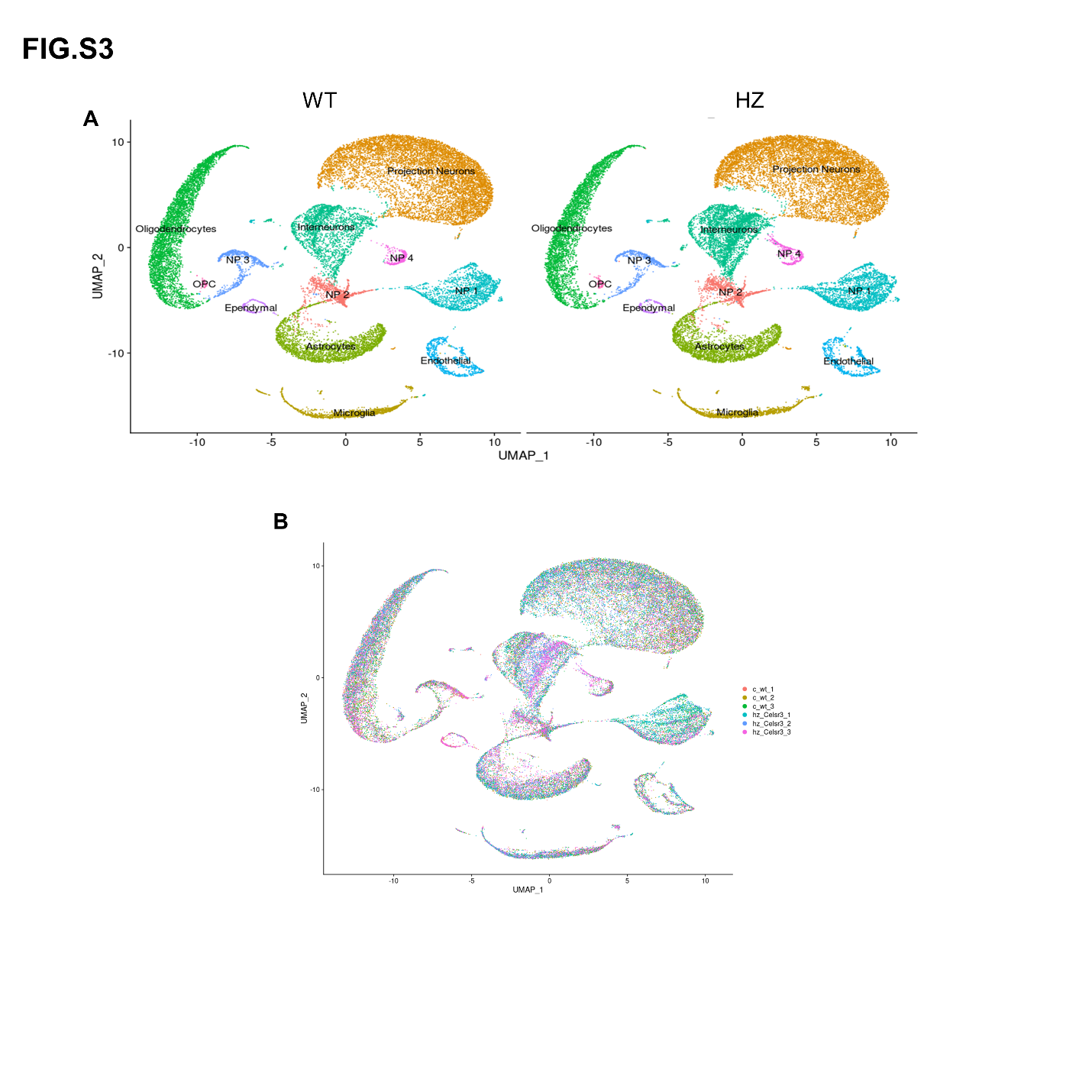
**Suppl. Fig. 9. Uniform Manifold Approximation and Projection (UMAP) of all cell types in wild-type (WT) and *Celsr3* heterozygous (HZ) mice.** (A) Uniform Manifold Approximation and Projection (UMAP) of all cell types including astrocytes, oligodendrocytes, microglia, projection neurons, interneurons, progenitor cells (OPC) and other neuronal populations (NP 1-4) for each genotype. (B) Overlay of the UMAP of all the individual samples used in this analysis.

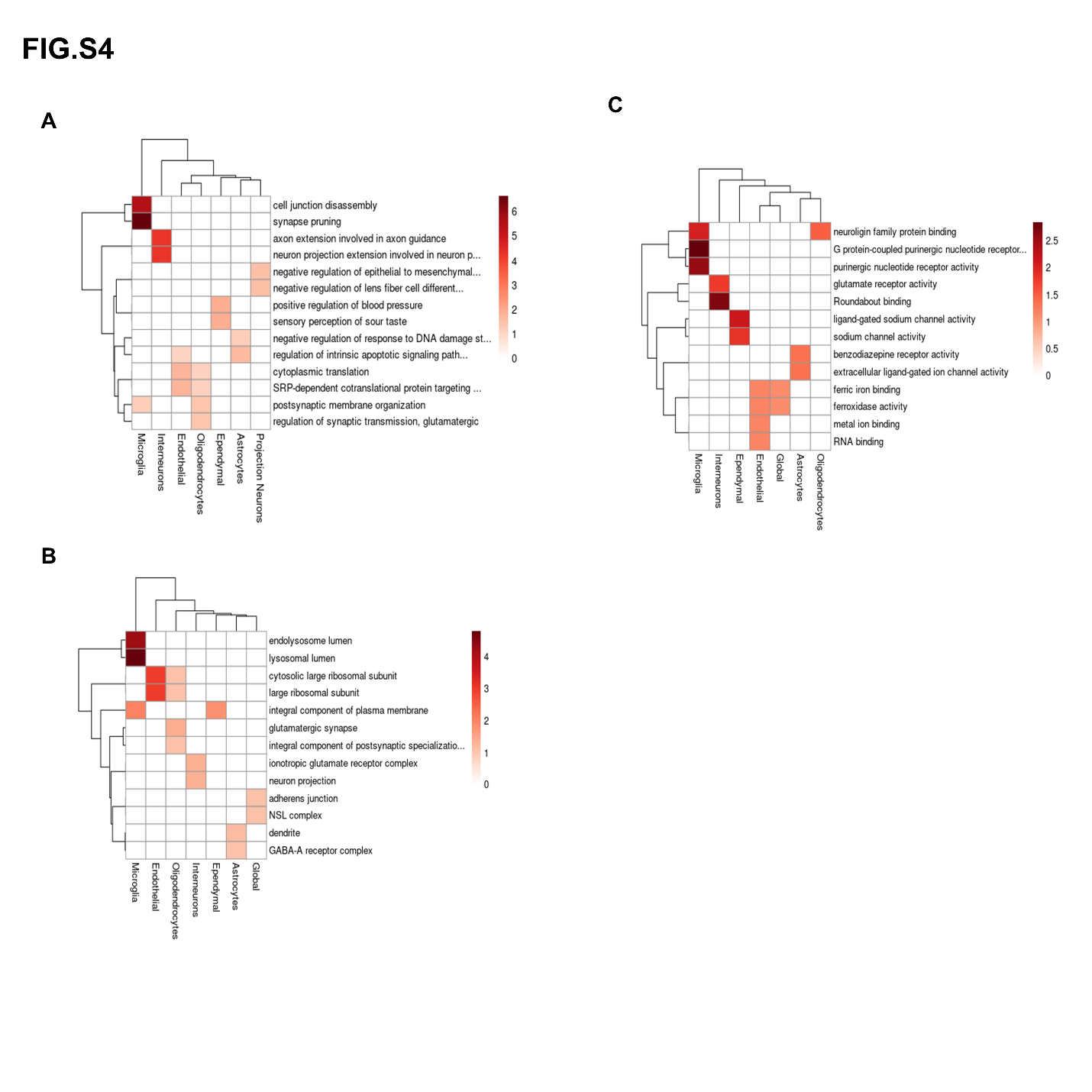

**Suppl. Fig. 10. Hierarchical clustering of top gene ontology (GO) terms across all cell populations in the overall comparison between wild-type (WT) and *Celsr3* heterozygous (HZ) mice.** A: GO Biological Process, B: GO Cellular Component; C: GO Molecular Function. For more details, see Suppl. Table 8 and text.

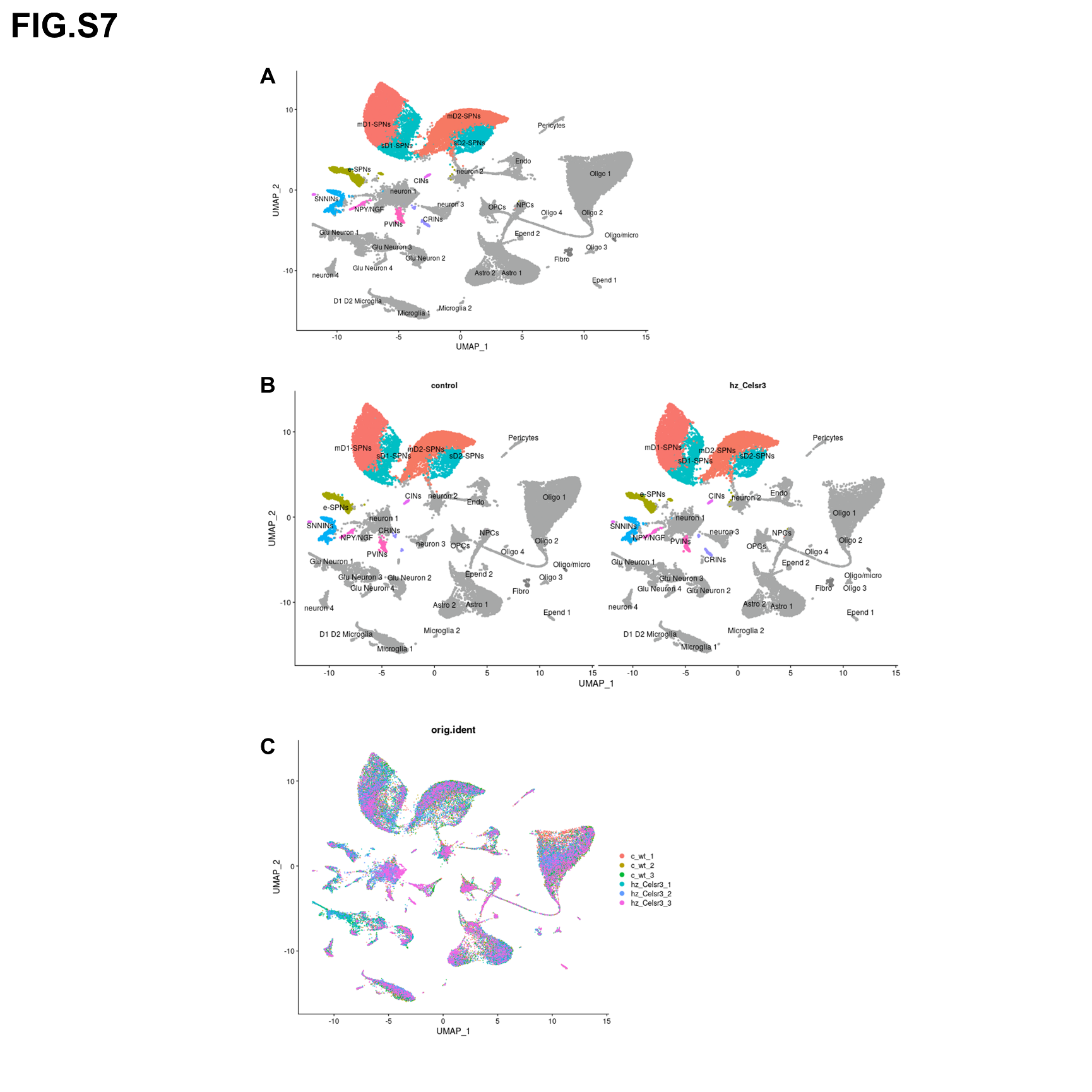

**Suppl. Fig. 11. Uniform Manifold Approximation and Projection (UMAP) of all cell types in wild-type (WT) and *Celsr3* heterozygous (HZ) mice.** (A) Uniform Manifold Approximation and Projection (UMAP) of all cell populations following SCTransformation with the generalized linear model, Gamma Poisson method. Cell types highlighted in the main manuscript are highlighted in color and all the others are displayed in grey. (B) UMAP of all cell populations for each genotype. (C) Overlay of the UMAP of all the individuals’ samples used in this analysis.

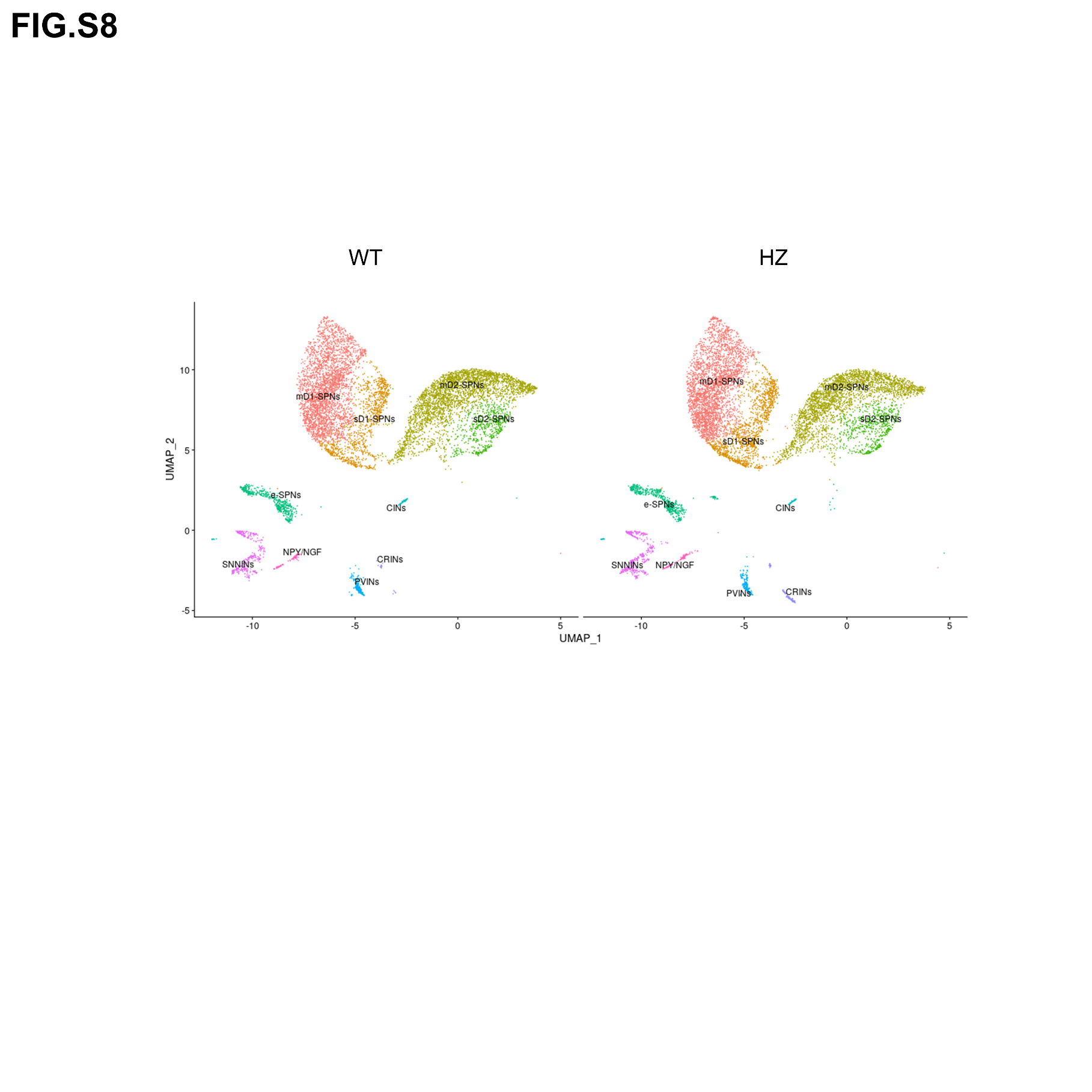

**Suppl. Fig. 12. Uniform Manifold Approximation and Projection (UMAP) of projection neurons and interneuron populations following SCTransformation with the generalized linear model, Gamma Poisson method, separated by genotype.** The most noticeable shifts are in the calretinin-positive interneurons. Abbreviations: mD1-SPNs, matrisome-associated, D_1_-positive striatal projection neurons; sD1-SPNs, striosome-associated, D_1_-positive striatal projection neurons; mD2-SPNs, matrisome-associated, D_2_-positive striatal projection neurons; sD2-SPNs, striosome-associated, D_2_-positive striatal projection neurons; e-SPNs, eccentric striatal projection neurons (exopatch neurons); CINs, cholinergic interneurons; SNNINs, somatostatin-NPY-NOS1-positive interneurons; NPY/NGF, NPY neurogliaform interneurons; PVINs, parvalbumin-positive interneurons; CRINs, calretinin-positive interneurons.

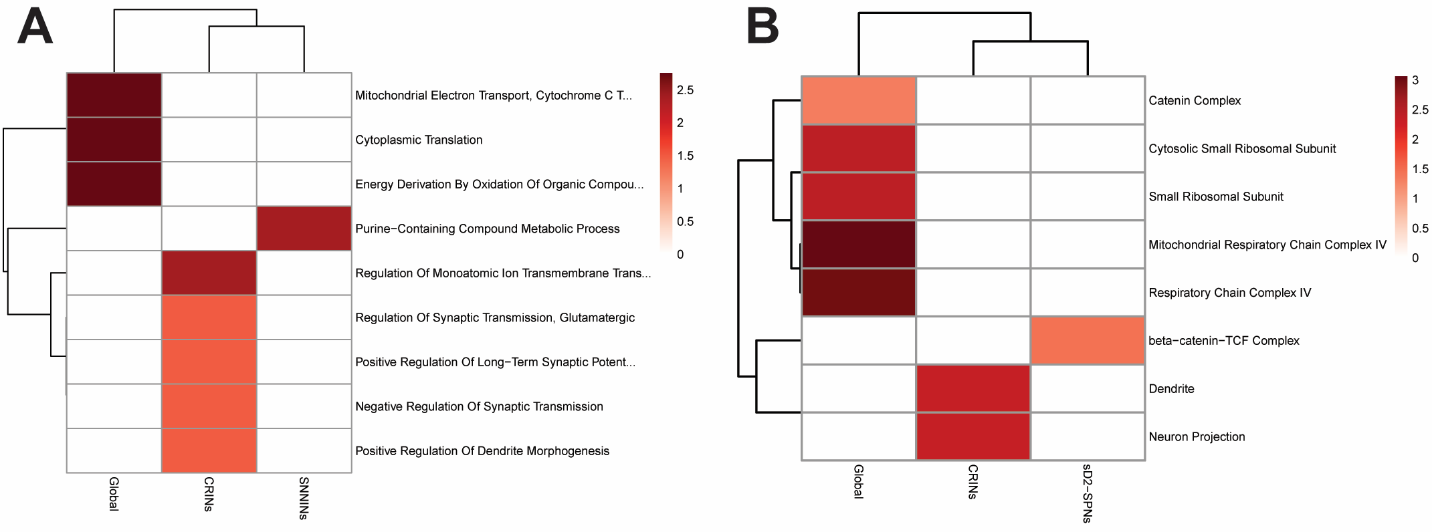
**Suppl. Fig. 13. Downregulated pathways in *Celsr3* heterozygous (HZ) mice, as outlined by hierarchical clustering of top gene ontology (GO) terms across all cell populations in comparison with wild-type (WT) mice.** A: GO Biological process, B: GO Cellular Component. For more details, see Suppl. Table 12 and text.

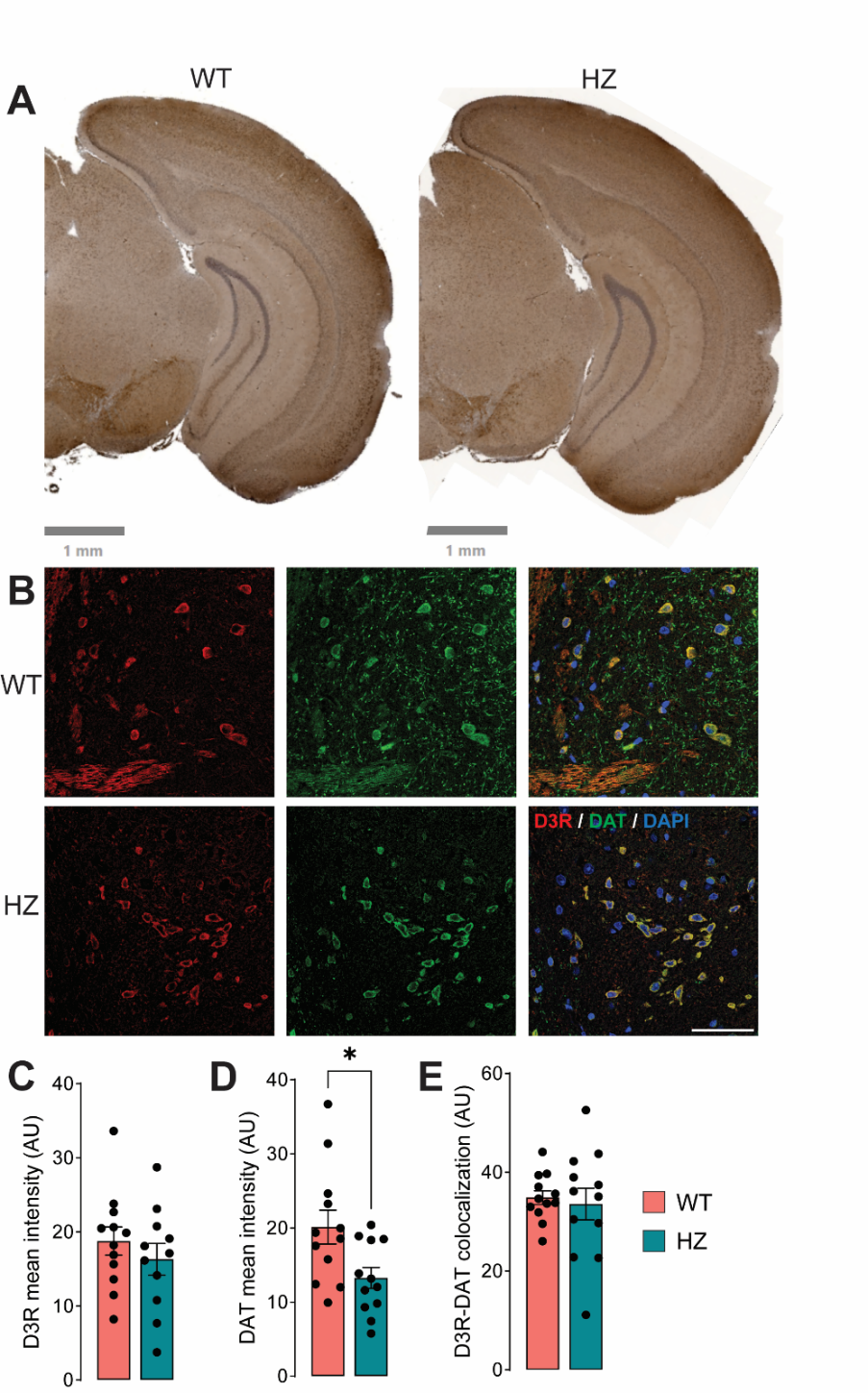

**Suppl. Fig. 14. Analyses of substantia nigra pars compacta of wild-type (WT) and *Celsr3* heterozygous (HZ) animals.** (A) Representative coronal sections of wild-type (WT) and heterozygous (HZ) animals showing immunohistochemical staining for dopamine transporter (DAT). The staining highlights the distribution of DAT in the striatum and cortex. Scale bar: 1 mm. (B) Immunofluorescent images showing D_3_ receptor (D3R, red) and dopamine transporter (DAT, green) co-localization in the striatum of WT and HZ animals. The merged images also include DAPI (blue) to label cell nuclei. Scale bar: 50 µm. (C-E) Quantification of D3R intensity (C), DAT intensity (D), and D3R-DAT colocalization (E) in WT and HZ animals. Semi-quantitative analyses were performed using ImageJ software (1.54i). Data are presented as mean ± SEM (n=3/group). *, p<0.05.

**SUPPLEMENTAL TABLES**

**Suppl. Table 1:** List of differentially expressed genes (DEGs) in bulk striatal tissue between *Celsr3* HZ and wild-type (WT) mice

**Suppl. Table 2:** List of Gene Ontology functional classes significantly enriched across differentially expressed genes (DEGs)

**Suppl. Table 3:** Data from spatial transcriptomic studies for all the clusters expressing *Drd1* and *Drd2* obtained from *Celsr3* HZ and wild-type (WT) mice highlighting the differentially expressed genes (DEGs) between clusters irrespective of the genotype.

**Suppl. Table 4:** Data from spatial transcriptomic studies for the selected striatal clusters depicting the list of differentially expressed genes (DEGs) in the comparison between *Celsr3* HZ and wild-type (WT) animals across all the different identified clusters and the GO analyses.

**Suppl. Table 5:** Data from spatial transcriptomic studies for the selected striatal clusters depicting the list of upregulated genes in the comparison between *Celsr3* HZ and wild-type (WT) animals across all the different identified clusters and the GO analyses.

**Suppl. Table 6:** Data from spatial transcriptomic studies for the selected striatal clusters depicting the list of downregulated genes in the comparison between *Celsr3* HZ and wild-type (WT) animals across all the different identified clusters and the GO analyses.

**Suppl. Table 7:** Data from single-nucleus (sn) transcriptomic studies for all the striatal cells obtained from *Celsr3* HZ and wild-type (WT) mice.

**Suppl. Table 8:** Data from single-nucleus (sn) transcriptomic studies depicting the list of differentially expressed genes (DEGs) across all the different identified clusters and the GO analyses.

**Suppl. Table 9:** Data from single-nucleus (sn) transcriptomic studies for the SPN and interneuron clusters obtained from *Celsr3* HZ and wild-type (WT) mice.

**Suppl. Table 10:** Data from single-nucleus (sn) transcriptomic studies depicting the list of differentially expressed genes (DEGs) across the SPN and interneuron clusters and the GO analyses.

**Suppl. Table 11:** Data from single-nucleus (sn) transcriptomic studies depicting the list of upregulated genes across the SPN and interneuron clusters and the GO analyses.

**Suppl. Table 12:** Data from single-nucleus (sn) transcriptomic studies depicting the list of downregulated genes across the SPN and interneuron clusters and the GO analyses.
